# Sugar Import Suppresses *Klebsiella pneumoniae* Mucoidy in cAMP-CRP-dependent Manner

**DOI:** 10.1101/2025.11.04.686645

**Authors:** Saroj Khadka, Gabriella Gates, Drew J. Stark, Bennett Allison, Laura A. Mike

## Abstract

*Klebsiella pneumoniae* causes over 700,000 global deaths annually and this number continues to rise due to increasing antimicrobial resistance. Hypervirulent *K. pneumoniae* (hv*Kp*) often causes severe invasive infections in community and hospital settings, typically originating from gut colonization. Capsular polysaccharide (CPS) and the associated features are a key hv*Kp* virulence factor. Altering CPS properties, such as mucoidy, in response to environmental cues enhances *K. pneumoniae* fitness. Although several physical and nutrient cues influence mucoidy, the molecular mechanisms by which host-relevant signals, such as sugars, regulate this key CPS property remain undefined. Here, we show that sugar import, not catabolism, broadly suppresses hv*Kp* mucoidy via cAMP-CRP signaling. Sugars suppress mucoidy by decreasing *rmpADC* promoter activity and the expression of the mucoidy regulator, *rmpD*. Moreover, this sugar-dependent regulation is conserved across multiple hv*Kp* strains. hv*Kp* associates with gut mucin and gut epithelial cells at a greater frequency when in a non-mucoid state. Together, our results suggest that sugar import broadly suppresses hv*Kp* mucoidy and increases bacterial association to host cells.

## 1. INTRODUCTION

*Klebsiella pneumoniae* is a Gram-negative bacterial pathogen responsible for a variety of nosocomial and community-acquired infections. It ranks as the fourth leading cause of bacterial infection-related deaths globally and accounts for an estimated 790,000 deaths annually.^1^ *K. pneumoniae* typically transmits via fecal-oral route and infects the lungs, bloodstream and urinary tract. Clinically important *K. pneumoniae* strains are broadly classified into two pathotypes: classical (c*Kp*) and hypervirulent (hv*Kp*). hv*Kp* particularly poses a public health threat due to its ability to cause severe invasive infections such as pyogenic liver abscess, meningitis and endophthalmitis in both healthy and immuno-compromised individuals. This is further exacerbated by the recent emergence of convergent isolates that are hypervirulent and antimicrobial resistant.^2,3^

Hypermucoviscosity (hereafter ‘mucoviscosity’ is referred to as ‘mucoidy’) is a defining virulence feature of hv*Kp*. Hypermucoid colonies exhibit a sticky phenotype linked to uniform capsular polysaccharide (CPS) chain length.^4,5^ Although its precise role in pathogenesis is not fully understood, *in vivo* studies support that hypermucoidy enhances virulence by impairing association to host immune cells like macrophages, ultimately promoting invasive infections.^6,7^ Even typically non-mucoid c*Kp* strains present increased virulence upon acquiring the hv*Kp* mucoidy regulator.^8^

In hv*Kp*, mucoidy is regulated by the *rmpADC* locus, which encodes three components: *rmpA*, an autoregulator; *rmpC,* a CPS biosynthesis regulator; and *rmpD*, the regulator of mucoid phenotype.^9^ RmpD increases mucoidy by directly interacting with Wzc, which controls CPS chain length during biosynthesis.^4^ Environmental factors such as pH, temperature, and iron and nutrient availability influence mucoidy through transcriptional and post-transcriptional regulation of *rmp* locus.^5,7,10–12^ In addition, genetic changes including spontaneous mutations in Wzc and phase-variation within the *rmpADC* locus further optimize hv*Kp* mucoidy.^5,13,14^ However, molecular mechanisms underlying several of these processes remain poorly defined, and additional regulatory signals are likely to be discovered.

Among these environmental factors, nutrient availability is a particularly important regulatory signal in the context of infection. Pathogens encounter distinct nutritional environments across different host niches. For example, glucose concentrations in the lung airway surface liquid (ASL) are 3-20 times lower than those in the plasma.^15,16^ Similarly, the human urinary tract maintains minimal free glucose concentration under normal conditions. However, disease states such as hyperglycemia and glycosuria elevate glucose level in these sites, which concomitantly promotes bacterial growth and persistence.^15,17,18^ The human gastrointestinal (GI) tract represents another nutritionally distinct environment that is both rich and diverse in available carbohydrates.^19^ The GI tract is lined with mucins that serves both as an innate immune defense and as a nutrient source to resident microbes.^20^ Mucins are structurally diverse glycoproteins, commonly composed of sugars like galactose, fucose, N-acetylglucosamine (GlcNAc), N-acetylgalactosamine (GalNAc) and N-acetylneuraminic acid (Neu5Ac). Microbial glycosidases and other carbohydrate-active enzymes liberate sugars from the dietary carbohydrates or host-derived mucins, providing energy sources to both commensal and pathogenic microbes.^21,22^ The role of sugars in host-pathogen interaction cannot be overstated as they modulate a wide array of virulence and fitness traits including capsule production, biofilm formation, antimicrobial resistance, secretion systems and motility.^23–27^ Collectively, sugars and other nutrients are important regulators of bacterial virulence and fitness. However, how sugar availability influences hv*Kp* mucoidy during gut colonization remains unknown.

*K. pneumoniae* extra-intestinal infections frequently originate from asymptomatic gut colonization. Up to 80% of hospitalized patients with *K. pneumoniae* gut carriage acquire subsequent infections from the colonizing strain.^28,29^ Thus, GI carriage is widely recognized as the primary reservoir and a key risk factor for *K. pneumoniae* infections.^30^ However, the factors that shift *K. pneumoniae* from colonizer to pathogen remain poorly understood. In particular, the influence of *K. pneumoniae* mucoidy regulation during gut colonization and subsequent dissemination to extra-intestinal sites remains unknown.

Our recent work demonstrated that arginine is a key environmental signal that promotes hv*Kp* mucoidy and niche-specific fitness.^7^ Interestingly, we also identified that certain sugars negatively modulate mucoidy, indicating that an opposing nutrient-based regulatory mechanism might exist in hv*Kp*.^7^ Given the carbohydrate-rich nature of the gut environment and the significance of GI carriage, we hypothesized that *K. pneumoniae* dynamically modulates mucoidy in response to available sugars to optimize its niche-specific fitness. To test this, we employed a genome-wide screen to identify bacterial genes and pathways involved in sugar-mediated mucoidy regulation.

Here, we demonstrated that host-relevant sugars suppressed hv*Kp* mucoidy and decreased CPS chain length uniformity by down-regulating *rmpD*. Mucoidy suppression required sugar import, not catabolism, and was observed across multiple hv*Kp* isolates.

In Gram-negative bacteria, sugar import is coupled to carbon catabolite regulation via cAMP-CRP. Disrupting cAMP-CRP signaling resulted in constitutively hypermucoid hv*Kp*, even in the presence of mucoidy-suppressing sugars. Finally, sugar-suppressed mucoidy increased bacterial binding to gut mucin and epithelial cells. Altogether, we present that sugar import generally suppresses mucoidy in a cAMP-CRP-dependent mechanism, suggesting that *K. pneumoniae* mucoidy is tailored *in vivo* by a balance between mucoidy-suppressing and -inducing nutrient signals.

## 2. RESULTS

### Sugars independently impact *K. pneumoniae* mucoidy and capsular polysaccharide abundance

To examine the impact of sugars on *K. pneumoniae* mucoidy, we selected the model hypervirulent strain KPPR1 and measured its mucoidy and CPS abundance using the standard sedimentation and uronic acid quantification assays in sugar-supplemented media.^31^ Specifically, KPPR1 was cultured in low-iron M9 minimal medium supplemented with 1% casamino acids (CAA) with or without 80 mM sugar (**M9+CAA+sugar**). Altogether, nine different sugars were tested: D-galactose (**Gal**), D-glucose (**Glc**), D-mannose (**Man**), L-rhamnose (**Rha**), L-arabinose (**Ara**), L-fucose (**Fuc**), D-xylose (**Xyl**), N-acetyl-D-glucosamine (**GlcNAc**) and N-acetyl-D-galactosamine (**GalNAc**). The sugars were selected if they fit one or more of the following parameters: known influence on mucoidy, metabolism by bacteria, and presence in host glycans and glycoproteins or bacterial CPS.^11,32–34^

Sugar supplementation (M9+CAA+sugar) significantly suppressed KPPR1 mucoidy, except for GalNAc, which had no significant effect (**Fig. 1A**). Notably, KPPR1 does not encode *agaVWEF,* a GalNAc phosphotransferase system (PTS) required for GalNAc transport in other Enterobacteriaceae.^35^ Meanwhile, sugar-specific transporters responsible for importing the other tested sugars are encoded in KPPR1. Furthermore, among the tested sugars, only GalNAc did not support KPPR1 growth when supplied as the sole carbon source, indicating that sugar-induced mucoidy is likely limited to consumable sugars (**Sup. Fig. 1I**). Sugar supplementation increases growth medium osmolarity and could exert osmotic stress on the bacterial membrane. As such, osmotic stress could suppress mucoidy. However, equimolar concentrations of GalNAc do not suppress mucoidy, indicating that the observed mucoidy modulation is not attributable to osmotic stress.

**Figure 1.**
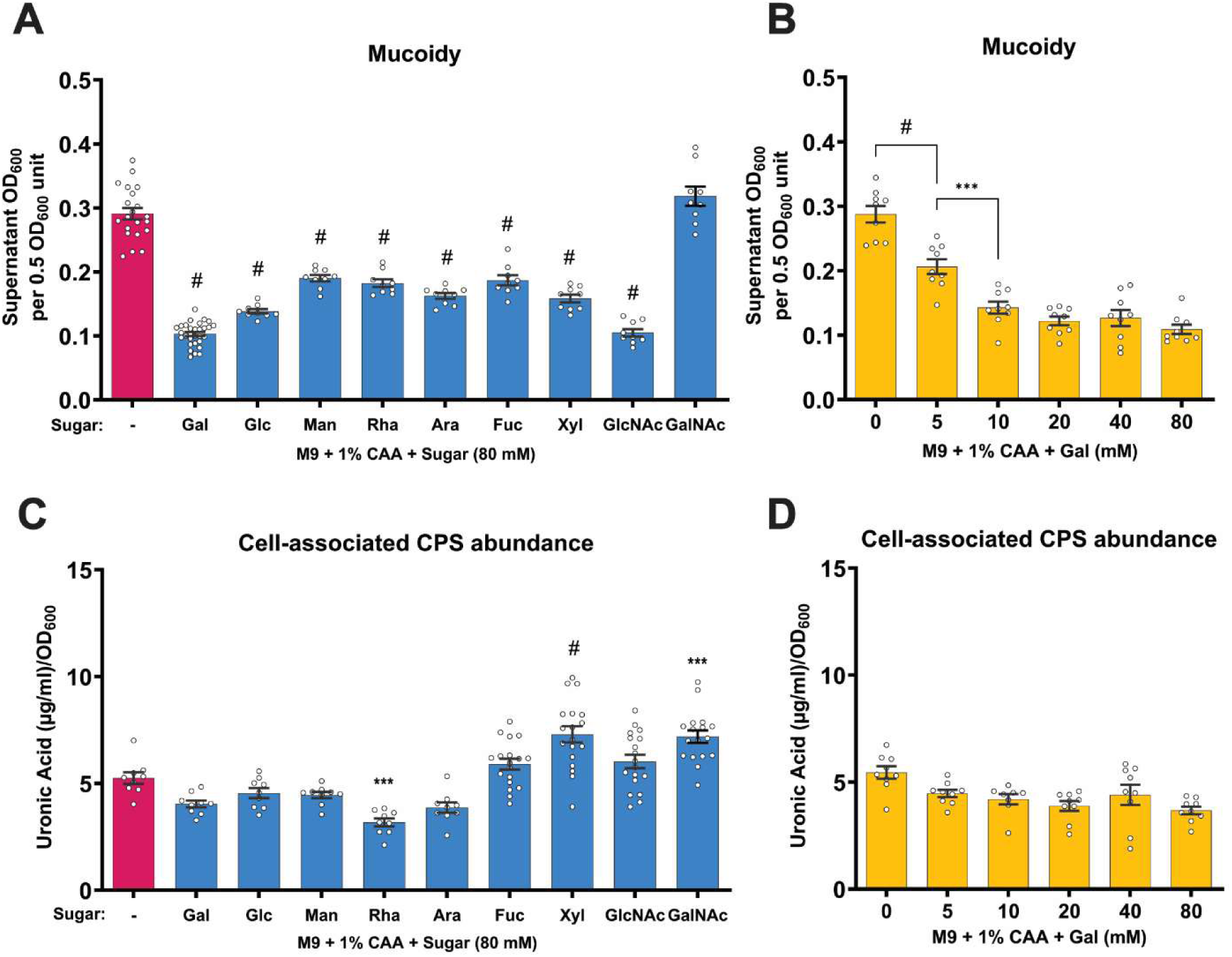
Sugar supplementation in minimal growth medium suppresses *K. pneumoniae* mucoidy independent of capsule abundance. *K. pneumoniae* strain KPPR1 was cultured in M9+CAA supplemented with **(A** and **C)** 80 mM of the respective sugar [D-galactose (Gal), D-glucose (Glc), D-Mannose (Man), L-rhamnose (Rha), L-arabinose (Ara), L-fucose (Fuc), D-xylose (Xyl), N-acetyl-D-glucosamine (GlcNAc) and N-acetyl-D-galactosamine (GalNAc)] or **(B** and **D**) varying concentrations of Gal ranging from 0 to 80 mM. **(A** and **B)** Mucoidy was determined by quantifying the supernatant OD_600_ after sedimenting 0.5 OD_600_ unit of culture at 1,000 x *g* for 5 mins. **(C** and **D)** Uronic acid abundance in crude CPS extracts was quantified for total CPS and supernatant CPS and normalized to OD_600_. Cell-associated CPS abundance was calculated by subtracting supernatant CPS from the total CPS content. Data presented are the mean, and error bars represent the standard error of the mean. Statistical significance was determined using one-way ANOVA with **(A** and **C)** Dunnett’s post-hoc test or **(B** and **D**) Šídák correction. Statistical significance was calculated by **(A** and **C)** comparing sugar-supplemented condition to sugar-deficient condition or **(B** and **D**) comparing adjacent pairs of bars. * *p* ≤ 0.05; ** *p* ≤ 0.01; *** *p* ≤ 0.001; # *p* ≤ 0.0001. Experiments were performed ≥3 independent times, in triplicate.

Although supplementing a variety of sugars at 80 mM broadly suppressed mucoidy, we wanted to determine whether the mucoidy suppression was concentration dependent. Therefore, we analyzed KPPR1 mucoidy at different concentrations of Gal. As low as 5 mM of Gal significantly suppressed KPPR1 mucoidy (**Fig. 1B**). It is possible that the mucoidy-suppressing effect of Gal at lower concentrations is counteracted by the mucoidy-inducing effect of arginine present in CAA.^7^

Hypermucoidy was conventionally attributed to increased CPS abundance, but recent studies have identified that increased CPS chain uniformity determines mucoidy.^5,6^ Consequently, CPS is essential for mucoidy. Thus, we also evaluated the cell-associated CPS content of KPPR1 under the same culture conditions (M9+CAA+sugar).^4,5^ Sugars had none to significant effects on KPPR1 CPS abundance depending on the sugar (**Fig. 1C**). Relative to M9+CAA, CPS abundance increased when GalNAc, Xyl, or GlcNAc was supplemented, but decreased when Gal, Rha, or Ara was supplemented (**Fig. 1C**). Notably, Gal did not significantly reduce KPPR1 CPS abundance in a dose-dependent manner (**Fig. 1D**). However, when either Gal or Glc were provided as the sole carbon source (**M9+Gal** and **M9+Glc**), KPPR1 mucoidy remained significantly suppressed, despite a nearly three-fold increase in CPS abundance compared to M9+CAA (**Sup. Fig. 2A-B**). In summary, we generally observed that consumable sugars decrease mucoidy, but have varied effects on cell-associated CPS abundance. We interpret these findings to indicate that sugars suppress *K. pneumoniae* mucoidy independently of their effect on CPS abundance.

### Sugar catabolism is not required to suppress *K. pneumoniae* mucoidy

Since *K. pneumoniae* mucoidy suppression was limited to consumable sugars, we investigated whether this regulatory process was linked to sugar metabolism or import. To test this, we provided two non-metabolizable sugar analogs, 2-deoxy-D-galactose (**2-D-Gal**) or 2-deoxy-D-glucose (**2DG**) in the M9+CAA medium and then measured KPPR1 mucoidy. These sugar analogs are imported via Gal- and Glc-specific transporters.^36,37^ However, neither 2-D-Gal nor 2DG is metabolized, and both sugars are unable to support KPPR1 growth as a sole carbon source (**Sup. Fig. 1A-B**). Surprisingly, both 2-D-Gal and 2DG suppressed KPPR1 mucoidy similar to Gal and Glc, despite being non-metabolizable sugar analogs **(Fig. 2A)**. Although 2-D-Gal supplementation significantly reduced CPS abundance, it remained unchanged following 2DG supplementation, re-affirming that sugar-suppressed mucoidy is not due to reduced CPS abundance (**Fig. 2B**). To further validate that sugar metabolism is not required for sugar-suppressed mucoidy, we ablated Gal catabolism by deleting the Leloir operon (*galETKM*). The Leloir pathway is required for the catabolism of Gal and as expected, the deletion mutant (***galETKM***) was unable to utilize Gal as a sole carbon source.^38^ Like WT, culturing *galETKM* in M9+CAA+Gal still significantly suppressed mucoidy compared to M9+CAA (**Fig. 2C**). These findings strongly support that Gal-mediated mucoidy suppression occurs upon sugar import and prior to catabolism.

**Figure 2.**
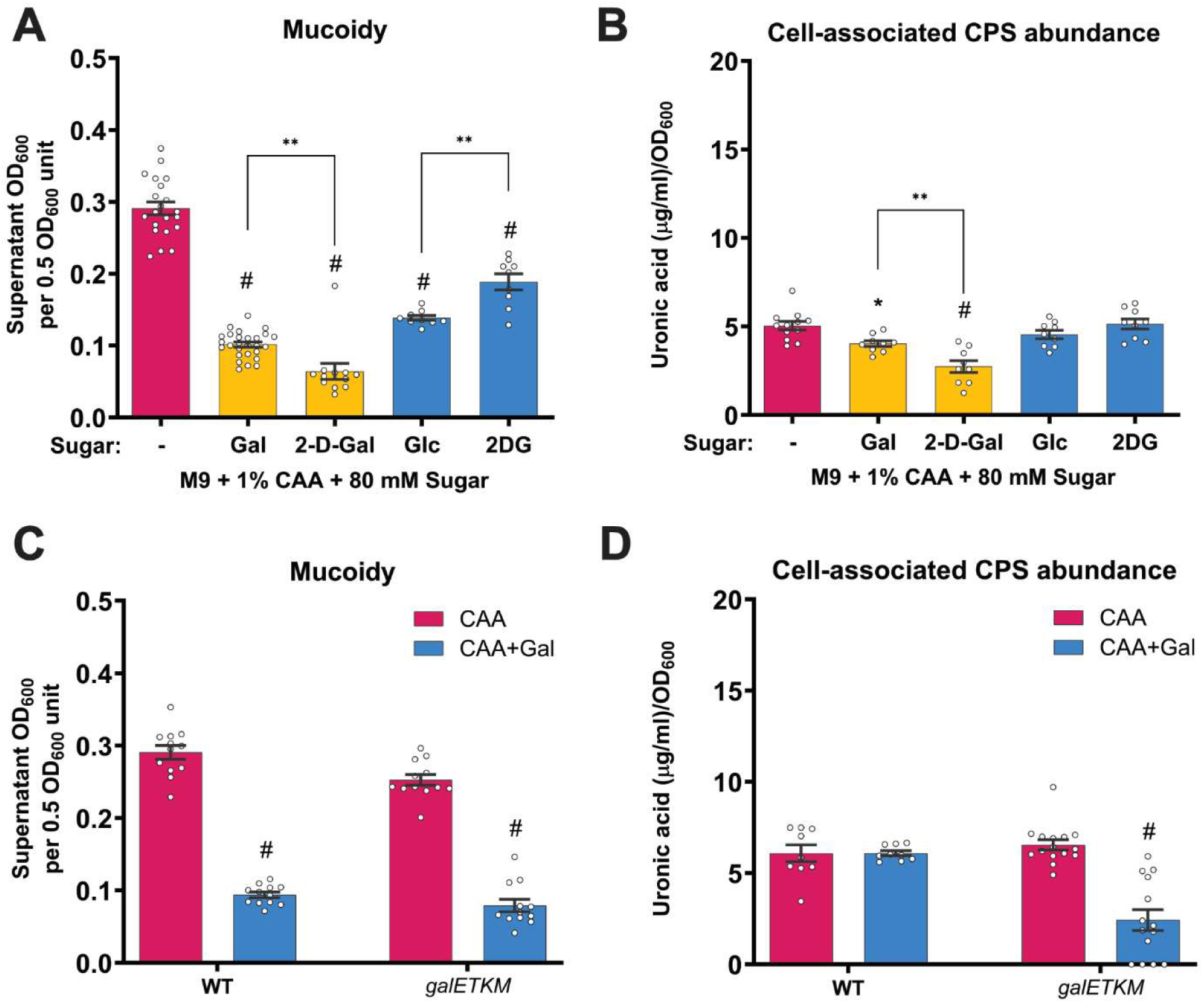
Sugar catabolism is dispensable for suppressing *K. pneumoniae* mucoidy. **(A-B)** KPPR1 was cultured in M9+CAA or M9+CAA+sugar [D-galactose (Gal), 2-deoxy-D-galactose (2-D-Gal), D-glucose (Glc), 2-deoxy-D-glucose (2DG)]. **(C-D)** KPPR1 wildtype (WT) and KPPR1 *galETKM::kan* were cultured in M9+CAA±Gal. **(A** and **C)** Mucoidy was determined by quantifying the supernatant OD_600_ after sedimenting 0.5 OD_600_ unit of culture at 1,000 x *g* for 5 mins. **(B** and **D)** Uronic acid abundance in crude CPS extracts was quantified for total CPS and supernatant CPS and normalized to OD_600_. Cell-associated CPS abundance was calculated by subtracting supernatant CPS from the total CPS content. Data presented are the mean, and error bars represent the standard error of the mean. Statistical significance was determined using **(A** and **B)** one-way ANOVA with Šídák correction and **(C** and **D)** two-way ANOVA with Šídák correction. Statistical significance was calculated by either comparing sugar-supplemented medium to the sugar-deficient medium or comparing adjacent pairs of bars. * *p* ≤ 0.05; ** *p* ≤ 0.01; # *p* ≤ 0.0001. Experiments were performed ≥3 independent times, in triplicate.

### Galactose downregulates the *rmpD* mucoidy regulator and decreases capsular polysaccharide chain length uniformity

In hv*Kp*, multiple regulatory inputs commonly act via transcriptional regulation of *rmpADC* locus making it the ‘central’ regulatory locus of mucoidy.^7,9,11,39,40^ Given the primary role of in regulating mucoidy, we hypothesized that sugars suppress mucoidy by transcriptionally down-regulating *rmpD*. To test this, we used a P*_rmp_*-*dasherGFP* fluorescent reporter construct to measure *rmpADC* promoter (P*_rmp_*) activity.^7^ We observed significantly reduced P*_rmp_* activity in M9+CAA+Gal compared to M9+CAA (**Fig. 3A**). The ∼3.3-fold decrease in promoter activity is consistent with the ∼3-fold decrease in mucoidy upon Gal supplementation (**Fig. 1A** and **3A**). Additionally, KPPR1 *rmpADC* transcript levels were significantly decreased by ∼5-fold in M9+CAA+Gal relative to M9+CAA (**Fig. 3B**). Together, these data suggest that Gal suppresses mucoidy by downregulating the mucoidy regulator, *rmpD*.

**Figure 3.**
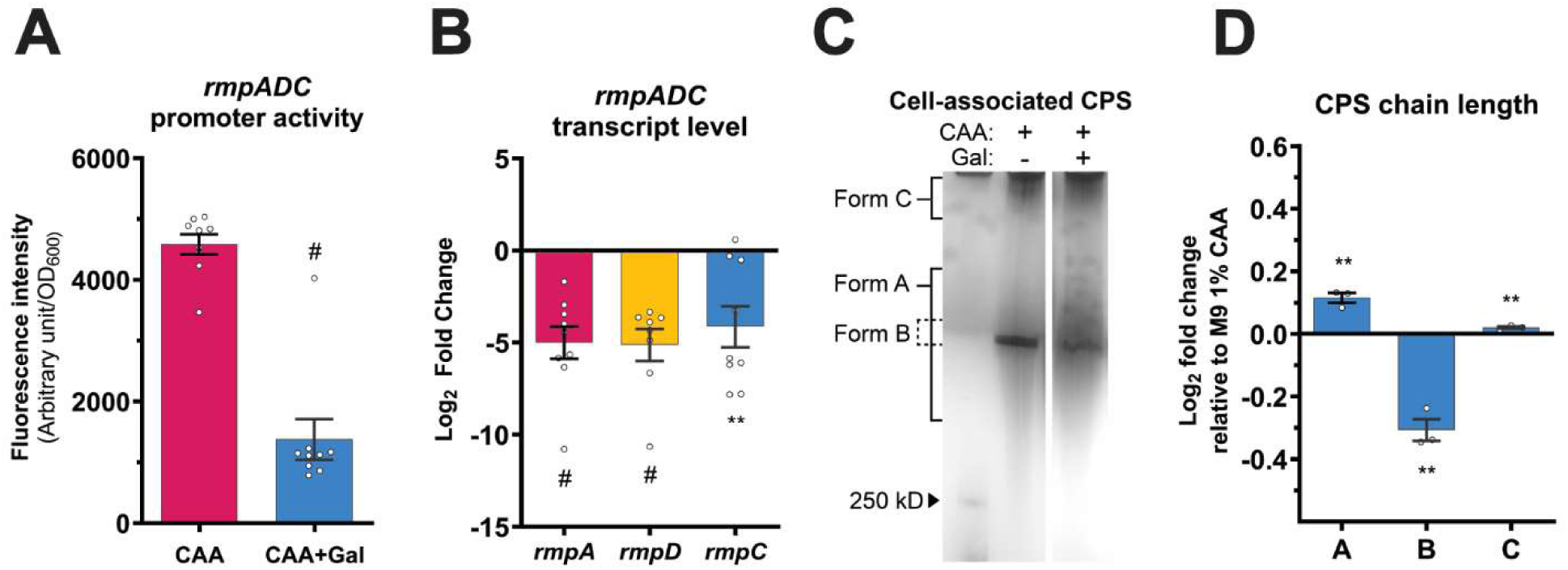
Galactose downregulates the regulator of mucoid phenotype (*rmpD*) and decreases CPS chain length uniformity. **(A)** KPPR1 carrying a GFP reporter vector (P*_rmp_*-*dasherGFP*) or **(B-D)** unmodified KPPR1 were cultured in M9+CAA±Gal. **(A)** KPPR1 *rmpADC* promoter (P*_rmp_*) activity was quantified by measuring the P*_rmp_*-driven GFP fluorescence intensity and normalized to OD_600_. **(B)** Relative transcript levels of KPPR1 *rmpA*, *rmpD* and *rmpC* were quantified by qRT-PCR in M9+CAA+Gal and compared to M9+CAA. **(C)** Cell-associated CPS was purified and resolved using a gradient SDS-PAGE and stained with 1% Alcian blue followed by silver stain. The presented gel image is representative of three independent experiments. **(D)** CPS chain length diversity was quantified by densitometric analysis using ImageJ. Data presented **(A, B** and **D)** are the mean, and the error bars represent the standard error of the mean. Statistical significance was determined using **(A)** unpaired t-test and **(B** and **D)** multiple unpaired t-tests with multiple-comparison correction using the Benjamini, Krieger and Yekutieli false discovery rate method. Statistical significance was calculated by comparing sugar-supplemented condition to sugar-deficient condition. ** *p* ≤ 0.01; # *p* ≤ 0.0001. Experiments were performed ≥3 independent times, in triplicate.

CPS chain length can be assessed by analyzing the migration patterns of purified CPS via SDS-PAGE.^41^ KPPR1 typically presents three distinct forms of CPS chains categorized by molecular weight (*i.e.* chain length) distribution. ‘Form A’ consists of CPS chains with diverse length (low- to high-molecular weight) and appears as a diffuse smear on a gel. ‘Form B’ represents high-molecular weight CPS chains of uniform chain length and appears as a distinct band. ‘Form C’ corresponds to ultra-high-molecular weight CPS chains retained close to loading wells. Hypermucoid strains predominantly synthesize high-molecular weight ‘form B’ CPS chains with uniform length distribution whereas low- and non-mucoid strains synthesize increased proportions of ‘form A’ CPS chains with diverse length.^4,5^ We purified cell-associated CPS from KPPR1 cultured in M9+CAA and M9+CAA+Gal, and analyzed the CPS chain length distributions by SDS-PAGE. Gal supplementation significantly increased the abundance of ‘form A’ CPS chains (diverse chain length) and decreased ‘form B’ chains (uniform chain length) compared to M9+CAA (**Fig. 3C-D**). Decreased CPS chain length uniformity (‘form B’) is consistent with Gal-suppressed mucoidy, as quantified by sedimentation assay (**Fig. 1A-B**).^5–7^ Collectively, our findings indicate that Gal import downregulates *rmpD* expression leading to the synthesis of CPS chains with decreased chain length uniformity and reduced mucoidy.

### Importable sugars downregulate *rmpADC* expression and broadly suppress mucoidy across hypervirulent *K. pneumoniae* strains

Since Gal-suppressed mucoidy was linked to *rmpD* downregulation, we next investigated whether other mucoidy-suppressing sugars similarly affected *rmpADC* expression. Using the P*_rmp_*-*dasherGFP* reporter, we measured *rmpADC* promoter activity in sugar-supplemented conditions. All tested sugars significantly reduced the P*_rmp_* activity, whereas GalNAc showed no impact on promoter activity, consistent with its lack of effect on mucoidy (**Fig. 4A** and **1A**). These findings indicate supplementation of other host-relevant sugars directly downregulates *rmpADC* expression, providing a mechanistic basis for sugar-suppressed mucoidy regulation.

**Figure 4.**
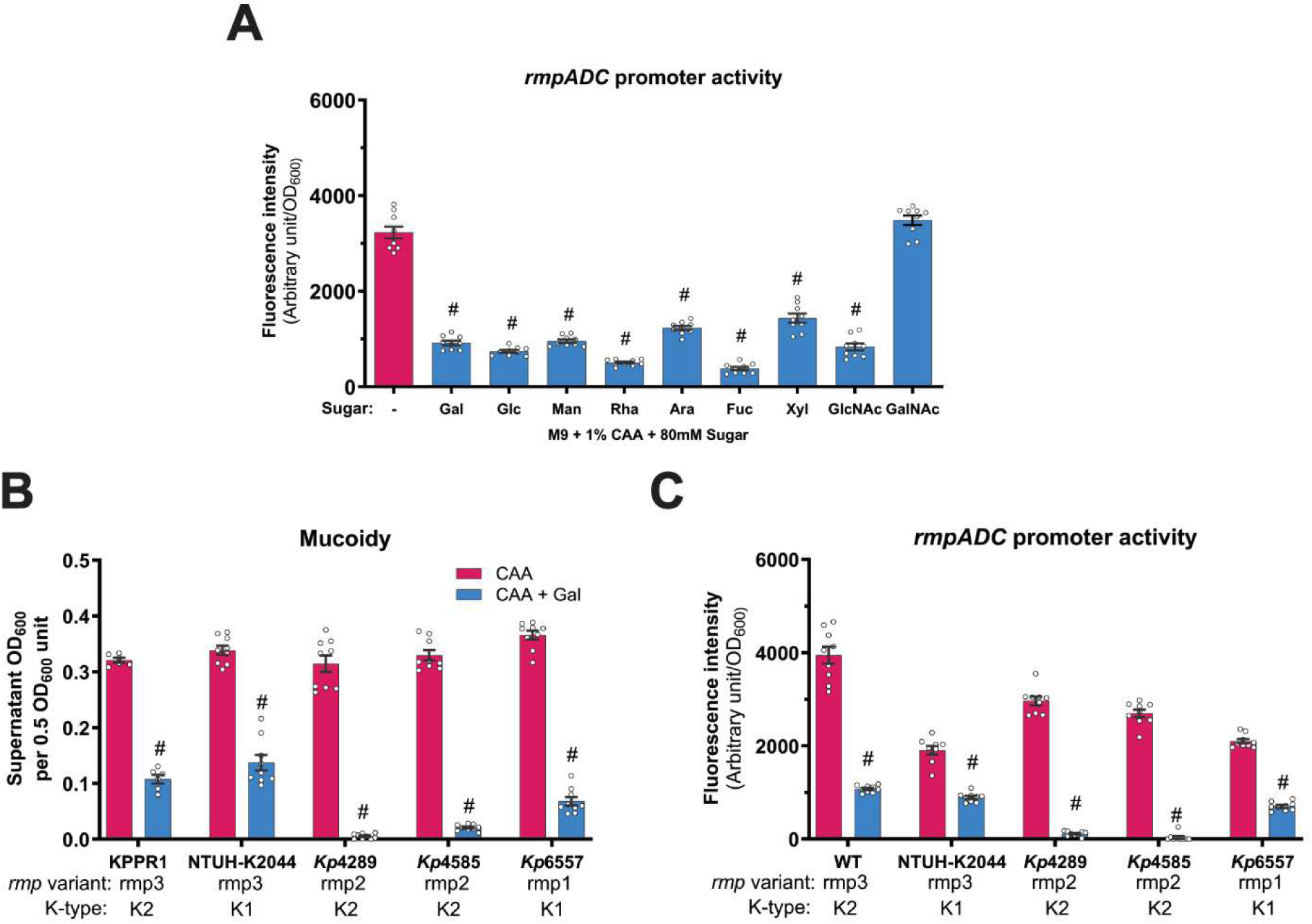
Sugar import reduces *rmpADC* promoter activity and galactose broadly suppresses mucoidy and *rmpADC* promoter activity in clinical hypervirulent *K. pneumoniae* strains. (A) KPPR1 P*_rmp_*-*dasherGFP* was cultured in M9+CAA+sugar and *rmpADC* promoter (P*_rmp_*) activity was quantified based on fluorescent protein expression normalized to OD_600_. **(B** and **C)** Additional hypervirulent *K. pneumoniae* strains were cultured in M9+CAA±Gal then mucoidy and P*_rmp_* activity were quantified. Laboratory strains tested were KPPR1 and NTUH-K2044, and clinical isolates were *Kp*4289, *Kp*4585 and *Kp*6557. **(B)** Mucoidy was determined by quantifying the supernatant OD_600_ after sedimenting 0.5 OD_600_ unit of culture at 1,000 x *g* for 5 mins. **(C)** P*_rmp_* activity of hypervirulent strains was quantified as in **(A)**. Data presented are the mean, and the error bars represent the standard error of the mean. Statistical significance was determined using **(A)** one-way ANOVA with Šídák correction and **(B** and **C)** two-way ANOVA with Šídák correction. Statistical significance was calculated by comparing sugar-supplemented condition to sugar-deficient condition. # *p* ≤ 0.0001. Experiments were performed ≥3 independent times, in triplicate.

To determine whether sugar-suppressed mucoidy is conserved across hv*Kp* strains, we tested the effect of Gal on additional hv*Kp* strains. We selected one laboratory hypervirulent strain (NTUH-K2044) and three clinical isolates (*Kp*4289, *Kp*4585 and *Kp*6557) in addition to KPPR1. Gal significantly suppressed mucoidy in all five hypervirulent strains (**Fig. 4B**). Furthermore, Gal significantly reduced P*_rmp_* activity in all tested hv*Kp* strains when quantified with the P*_rmp_*-dasherGFP reporter (**Fig. 4C**).^7^ This suggests that the import of Gal, and likely other sugars, generally suppress mucoidy by transcriptionally downregulating *rmpADC* via a regulatory mechanism conserved across hypervirulent strains.

### Transposon screen identifies genes associated with sugar-suppressed mucoidy

To identify bacterial factors involved in sugar-suppressed mucoidy, we screened a *K. pneumoniae* transposon library for mutants demonstrating increased mucoidy in Gal. We used a previously published, ordered KPPR1 transposon library consisting of 3,733 mutants, representing approximately 72% of all open reading frames (ORFs).^5,42^ A high-throughput sedimentation assay adapted for a 96-well plate was employed in two successive rounds to screen for mutants with increased mucoidy in M9+CAA+Gal. In the primary screen, 261 mutants were identified as preliminary hits which were refined to 40 mutants following the secondary screen in M9+CAA+Gal (**Sup. Fig. 3**). The hits were validated using a standard, full-scale (3 mL) sedimentation assay, assessing the mutants for increased mucoidy in M9+CAA+Gal compared to WT.^41^ Ultimately, *n = 15* transposon mutants were confirmed as final hits that had increased mucoidy in M9+CAA+Gal (**Fig. 5A**). Targeted gene deletion mutants of each of the 15 hits were generated and their mucoidy was measured. All but three mutants (*rpoN*, *aroK* and *yceG*) maintained increased mucoidy in M9+CAA+Gal (**Fig. 5B**). Whole genome sequencing revealed no secondary mutations that could explain the discrepancy in mucoidy between the transposon and deletion mutants of these three genes. Overall, the 15 identified gene hits spanned diverse cellular processes. Most were associated with carbon-responsive global regulation (*cyaA*, *crp*, *csrD*) and nitrogen regulation and signaling (*ntrB*, *ntrC*, *ptsN*, *rpoN*) (**Fig. 5A—B**). Additional hits included genes implicated in antimicrobial peptide transport (*sapA* and *sapB*), inorganic phosphate uptake (*pitA*), cell-wall remodeling (*yceG*), cofactor biosynthesis (*epd*), pentose phosphate pathway (*rpe*) and amino acid biosynthesis (*aroK*) (**Fig. 5A—B**).

**Figure 5.**
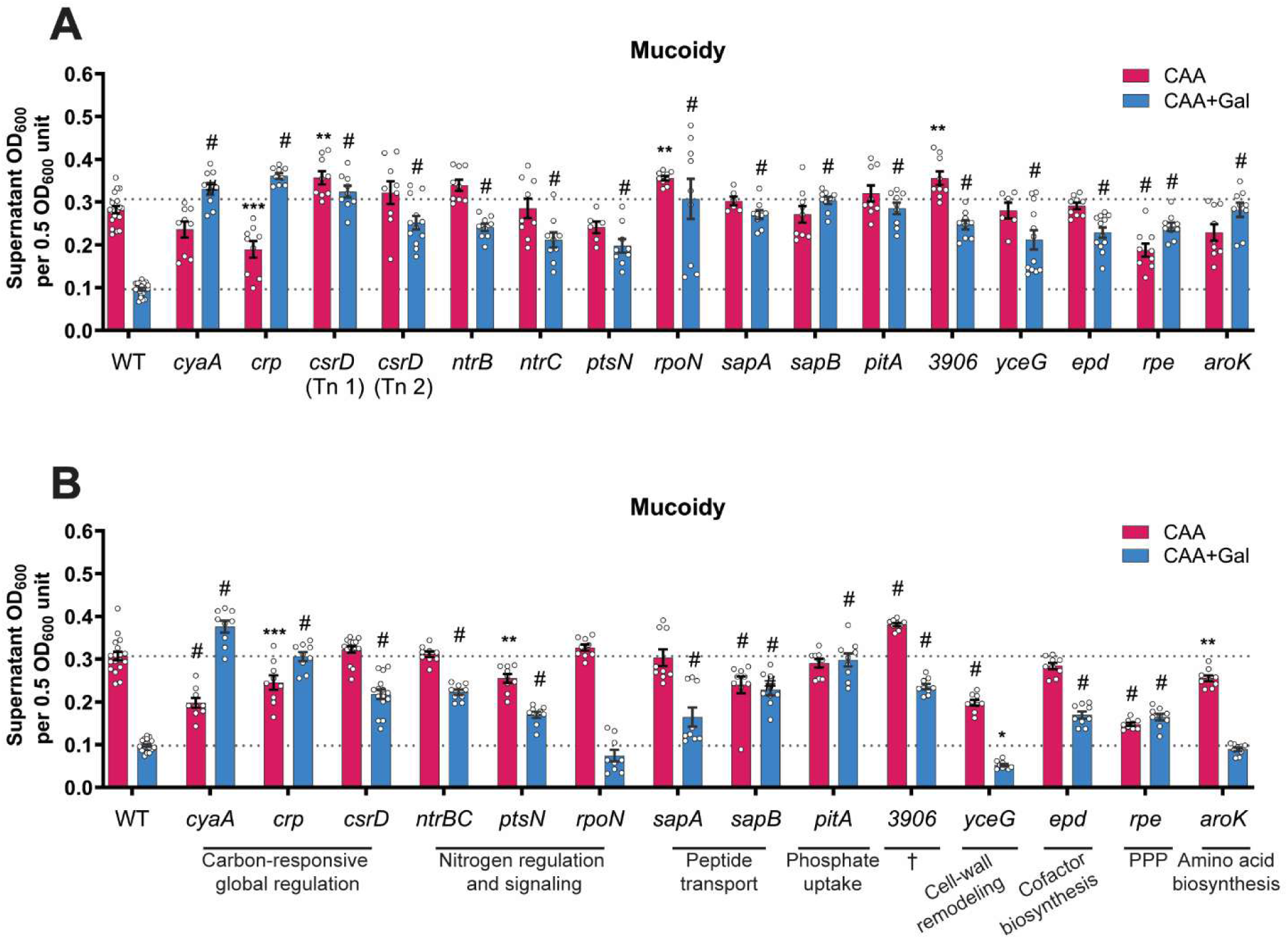
Transposon screen identifies *K. pneumoniae* mutants with increased mucoidy in galactose-supplemented medium. **(A)** Transposon (Tn) mutant hits were cultured in M9+CAA±Gal and validated for increased mucoidy in M9+CAA+Gal compared to M9+CAA by quantifying the supernatant OD_600_ after centrifugation at 1,000 x *g* for 5 mins. **(B)** Isogenic gene deletion mutants were generated for each KPPR1 Tn mutant identified as positive hits in **(A)**. The mutants were cultured in M9+CAA±Gal, and their mucoidy was determined by quantifying the supernatant OD_600_ after centrifugation at 1,000 x *g* for 5 mins. Data presented are the mean, and the error bars represent the standard error of the mean. Statistical significance was determined using two-way ANOVA with Dunnett’s post-hoc test. Statistical significance was calculated by comparing each mutant to the WT in the respective growth medium. * *p* ≤ 0.05; ** *p* ≤ 0.01; *** *p* ≤ 0.001; # *p* ≤ 0.0001. Experiments were performed ≥3 independent times, in triplicate. † Gene of unknown function; PPP: pentose phosphate pathway.

### A global regulator, cAMP receptor protein (CRP), is required for sugar-suppressed mucoidy

CRP is a global carbon catabolite regulator known to play a role in virulence regulation in *Klebsiella* and other bacterial species.^43–46^ CRP forms a complex with cAMP, the adenylate cyclase (CyaA) product, to upregulate or downregulate various metabolic and virulence-associated genes. cAMP synthesis requires CyaA activation, which is primarily regulated by sugar import through sugar-specific phosphotransferase (PTS) systems. Given these established functions of CRP and the identification of both *cyaA* and *crp* as major transposon hits in our screen, we further investigated the role of cAMP-CRP in sugar-mediated mucoidy regulation. We measured mucoidy of a KPPR1 *cyaA* and *crp* deletion mutants cultured in M9+CAA with or without Gal. As expected, both the mutants lacked intracellular cAMP and restored mucoidy in M9+CAA+Gal (**Fig. 6A** and **C**). Meanwhile, expressing *cyaA* and *crp in trans* using native promoters restored Gal-suppressed mucoidy to WT levels in M9+CAA+Gal (**Sup. Fig. 4A**). Furthermore, Gal supplementation increased P*_rmp_* activity more than ∼9.5-fold in both *cyaA* and *crp* mutants (**Fig. 6D**). Interestingly, in M9+CAA there was no significant change in P*_rmp_* activity (**Fig. 6D**), yet both mutants had significant decreases in mucoidy compared to WT (**Fig. 6A**). The increased mucoidy and P*_rmp_* activity of *cyaA* and *crp* in M9+CAA+Gal correlated with an increased abundance of uniform-length CPS chains (‘Form B’) relative to WT (**Fig. 6E-F**). In contrast, CPS chains with diverse lengths, represented by ‘Form A’, decreased in abundance (**Fig. 6E-F**). Collectively, these observations implicate cAMP-CRP signaling in sugar-dependent

**Figure 6.**
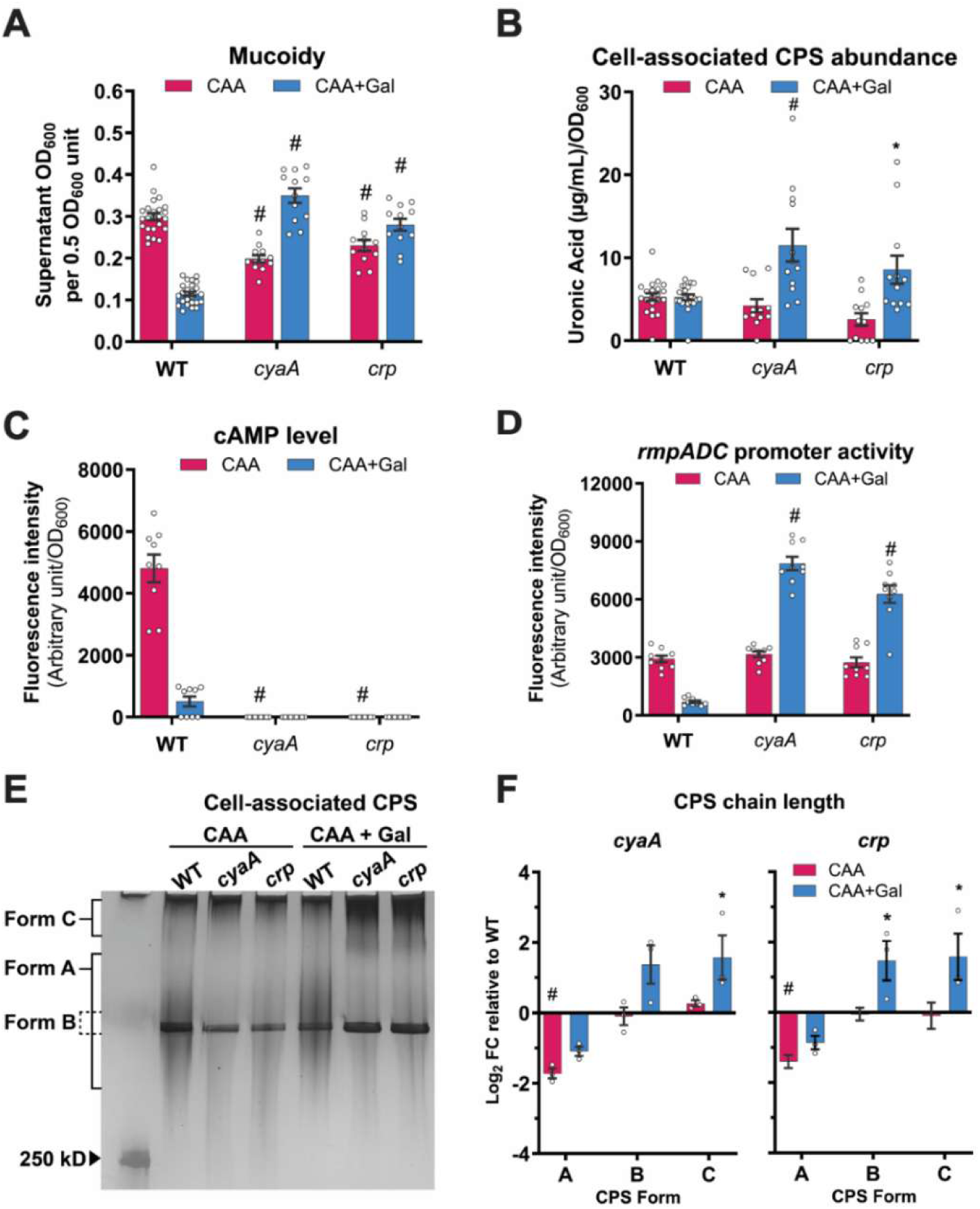
Sugar-suppressed mucoidy is linked to the cAMP-CRP global regulator. KPPR1 wildtype (WT), and *cyaA* and *crp* mutants were cultured in M9+CAA±Gal. **(A)** Mucoidy was determined by quantifying the supernatant OD_600_ after centrifugation at 1,000 x *g* for 5 mins. **(B)** Cell-associated CPS was extracted and measured for uronic acid content. **(C)** Intracellular cAMP levels and **(D)** P*_rmp_* activity were measured using a P*_rmp_* and cAMP-CRP-responsive dual fluorescent reporter. Cell-associated CPS were **(E)** resolved by SDS-PAGE and **(F)** analyzed for chain length diversity by densitometric analysis in ImageJ. Data presented are the mean, and the error bars represent the standard error of the mean. Statistical significance was determined using two-way ANOVA with Dunnett’s post-hoc test where each mutant was compared to WT in the same growth medium. * *p* ≤ 0.05; # *p* ≤ 0.0001. CPS staining was performed independently three times with a cumulative total of *n = 3* biological replicates. All other experiments were performed ≥3 independent times, in triplicate.

### *K. pneumoniae* mucoidy regulation

Next, we asked whether cAMP-CRP broadly drives sugar-suppressed mucoidy. We quantified intracellular cAMP levels in KPPR1 cultured in minimal medium supplemented with a panel of sugars. Analogous to Gal, each imported sugar significantly reduced intracellular cAMP levels (**Fig. 7A**). Conversely, the *cyaA* mutant had significantly increased mucoidy in M9+CAA supplemented with each sugar (**Fig. 7B**). Altogether, our data strongly suggests that the general regulatory effect of sugars on *K. pneumoniae* mucoidy is mediated by sugar import-driven changes in intracellular cAMP.

**Figure 7.**
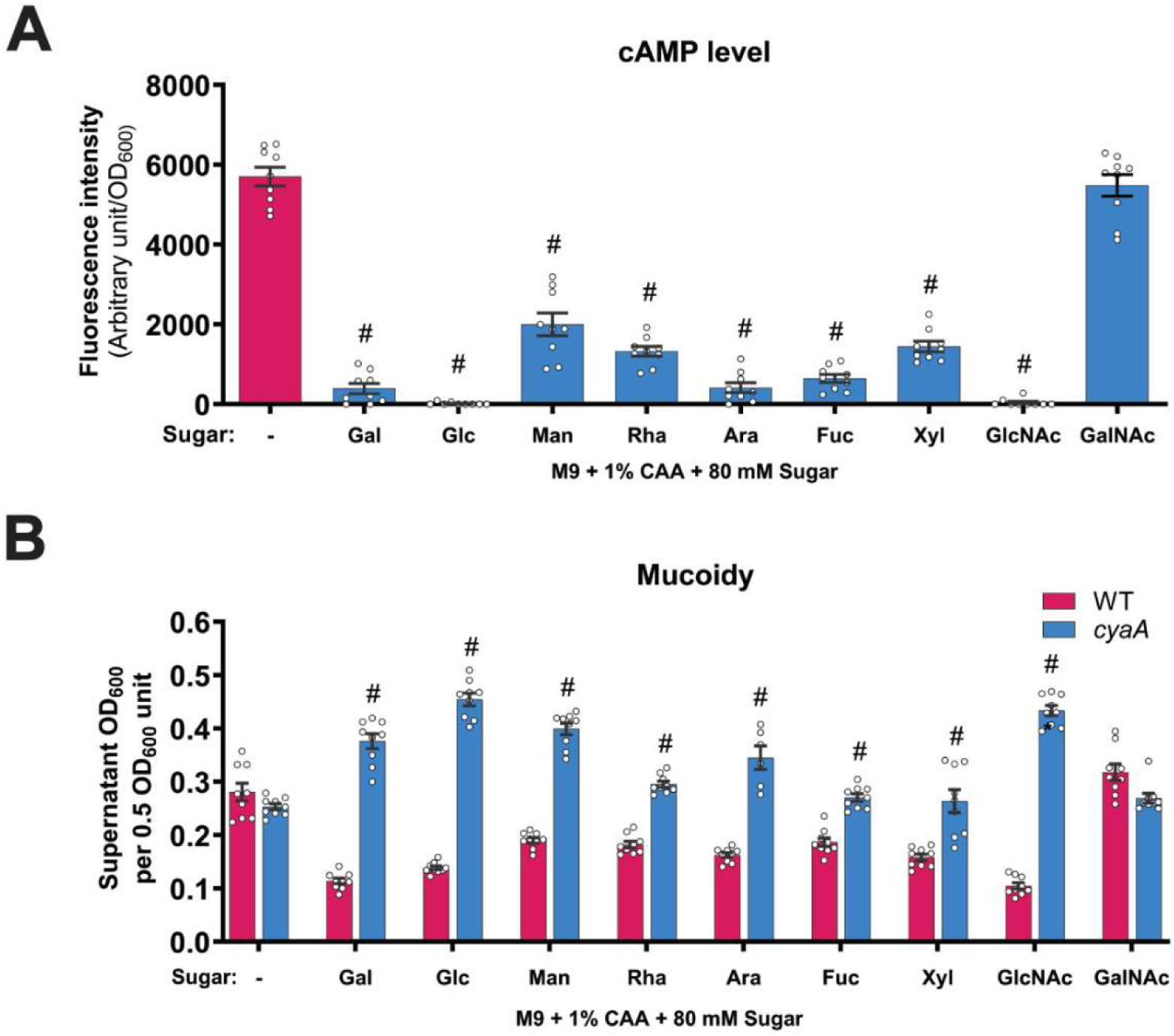
Sugar import broadly reduces intracellular cAMP and suppresses mucoidy via the cAMP-CRP regulatory pathway. KPPR1 wildtype (WT) and/or *cyaA* mutant were cultured in M9+CAA±sugar. **(A)** Intracellular cAMP levels of WT cultured in different sugars were measured using a cAMP-CRP-responsive fluorescent reporter and normalized to OD_600_. **(B)** The mucoidy of WT and *cyaA* cultured in different sugars was determined by quantifying the supernatant OD_600_ after centrifugation at 1,000 x *g* for 5 mins. Data presented are the mean, and the error bars represent the standard error of the mean. Statistical significance was determined using **(A)** one-way ANOVA with Šídák correction and **(B)** two-way ANOVA with Šídák correction by comparing **(A)** sugar-added condition to M9+CAA or **(B)** mutant to WT. # *p* ≤ 0.0001. All experiments were performed ≥3 independent times, in triplicate.

### Non-mucoid *K. pneumoniae* associate at higher frequency with gut mucin and epithelial cells

*K. pneumoniae* is able to invade and translocate across gut epithelial cells via a transcellular route.^47^ Moreover, low- and non-mucoid *K. pneumoniae* strains typically associate more effectively than hypermucoid strains to host cells, including lung epithelial cells.^42,48^ However, pathogens must first overcome the mucus barrier to reach the epithelial (primarily composed of mucin), which limits access to the epithelial surface. We therefore examined whether mucoidy suppression impacts *K. pneumoniae* interaction with gut mucin and epithelial cells, thereby affecting key steps in gut colonization and eventual dissemination.

To test the effect of mucoidy suppression on bacterial mucin binding, we performed a solid-phase mucin binding assay using semi-purified porcine gastric mucin (PGM). A significantly higher proportion (∼5-fold) of KPPR1 WT pre-grown in M9+CAA+Gal were bound to PGM compared to WT grown in M9+CAA **(Fig. 8A)**. Furthermore, a constitutively non-mucoid *rmpD* mutant, bound to mucin significantly higher (∼7.5-fold) than WT in CAA, regardless of Gal supplementation **(Fig. 8A)**. These findings suggest that modulation of mucoidy by host-derived sugars may influence interactions between *K. pneumoniae* and mucin.

**Figure 8.**
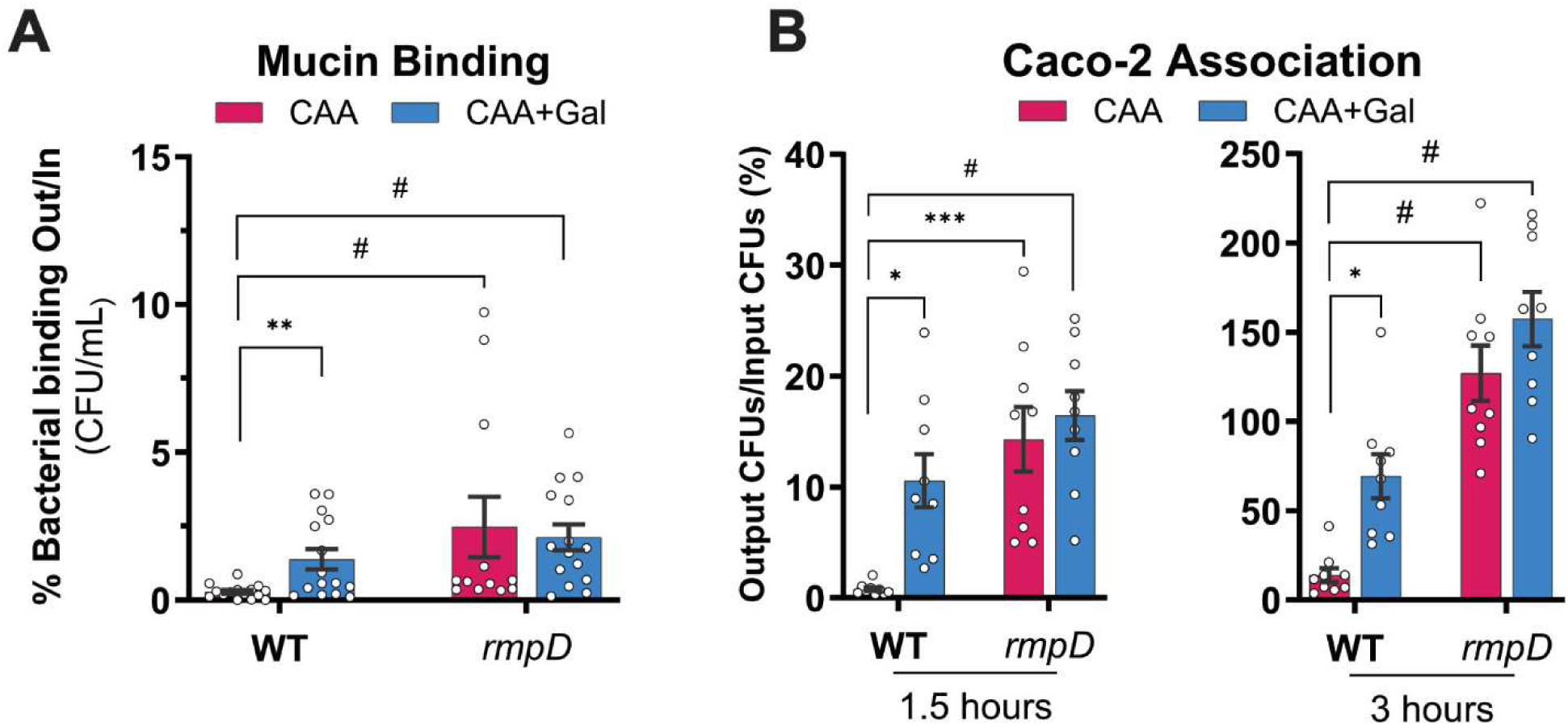
Non-mucoid *K. pneumoniae* bind porcine gastric mucin and gut epithelial cells more tightly. KPPR1 WT and *rmpD* pre-cultured in M9+CAA±Gal binding to porcine gastric mucin (PGM) and infection rate of human immortalized intestinal epithelial cells (Caco-2) was quantified. **(A)** A Nunc MaxiSorp plate was coated with 0.15 mg of crude PGM and treated with ∼10^5^ KPPR1 WT and *rmpD* cells. After treatment, bacterial binding was measured by enumerating colony forming units (CFU) relative to input bacterial load. **(B)** A monolayer of Caco-2 cells was infected with KPPR1 WT or *rmpD* for 1.5 and 3 hours with an MOI of 10. Infected cells were washed with PBS and then Caco-2 cells were lysed with Triton X-100. Total bacterial association was quantified by enumerating CFU relative to input bacterial load. Data presented are the mean, and the error bars represent standard error of the mean. Statistical significance was determined using (A) Mann-Whitney test with multiple-comparison correction using the Benjamini, Krieger and Yekutieli false discovery rate method, and (B) two-way ANOVA with Šídák’s multiple comparison test. * *p* ≤ 0.05; ** *p* ≤ 0.01; *** *p* ≤ 0.001; # *p* ≤ 0.0001. Experiments were performed ≥3 independent times, in triplicate.

We next assessed whether mucoidy suppression alters *K. pneumoniae* adherence and invasion of Caco-2 intestinal epithelial cells. KPPR1 WT grown in M9+CAA+Gal associated with Caco-2 cells more efficiently at 1.5 hours (12-fold) and 3 hours (>4-fold) compared to WT in M9+CAA **(Fig. 7B)**. Similarly, WT in M9+CAA+Gal invaded Caco-2 cells at a 3-fold higher rate than WT in M0+CAA after 1.5 hours, which was reduced to a modest 1-fold higher rate at 3 hours **(Fig. 7C)**. The constitutively non-mucoid *rmpD* mutant consistently showed higher association with Caco-2 cells compared to WT, irrespective of Gal (**Fig. 7B-C**). Together, these results reinforce that sugars suppress mucoidy through an *rmpD*-dependent mechanism that enhances *K. pneumoniae* interaction with gut mucin and epithelial cells, with potential consequences in gut colonization and dissemination.

## 3. DISCUSSION

Nutrient signals like amino acids and sugars are known modulators of *K. pneumoniae* mucoidy, but the mechanism by which sugars influence mucoidy has remained undefined. Here, we investigated how individual sugars influence transcriptional responses and CPS properties that dictate mucoidy. We found that sugars broadly suppress *K. pneumoniae* mucoidy by downregulating *rmpADC* and diversifying CPS chain length. This regulation is coupled with sugar import, requiring cAMP-CRP, and is independent of sugar catabolism. Moreover, sugar-suppressed mucoidy is conserved across hypervirulent *K. pneumoniae* strains and could influence host-pathogen interactions in the gut and other niches. These data support and expand our understanding of how *K. pneumoniae* metabolic status regulates the mucoid state. We propose a model whereby sugars and amino acids exert opposing effects on *K. pneumoniae* CPS properties, thereby integrating nutritional cues to fine-tune virulence and fitness within host microenvironments.

We systematically tested eleven individual sugars for their effects on *K. pneumoniae* mucoidy. All, except GalNAc, suppressed KPPR1 mucoidy (**Fig. 1A**). Notably, GalNAc was the only sugar unable to support growth as a sole carbon source, which is consistent with the absence of GalNAc transporter (*ageVWEF*) in KPPR1 (**Sup. Fig. 1**). These data suggest that consumable sugars broadly suppress *K. pneumoniae* mucoidy. However, assays using non-metabolizable sugar analogs (2-d-Gal and 2DG) and a galactose catabolic mutant (*galETKM*) revealed that mucoidy suppression is triggered by sugar import, not catabolism (**Fig. 2A** and **2C**). Our data support that sugar-suppressed mucoidy is independent of osmotic stress caused by sugars or pH changes due to sugar catabolism. Specifically, equimolar GalNAc does not suppress mucoidy, indicating that sugar-suppressed mucoidy is not likely a bacterial response to osmotic stress. Additionally, because *galETKM* mutant cannot metabolize Gal, and WT cannot metabolize 2-d-Gal, the medium pH remains stable, yet we still observe sugar-suppressed mucoidy in these conditions (**Sup. Fig. 5**). To the best of our knowledge, this is the first study to extensively investigate the relationship between sugar availability and mucoidy. Similar to our experimental approach, a previous study examined mucoidy in CAA supplemented with glucose, glycerol or fucose.^11^ While this study did not specifically compare mucoidy in sugar-added conditions to M9+CAA, the mucoidy was lower relative to LB, a growth medium that elicits similar mucoidy levels as M9+CAA in our experience.

Although mucoidy and CPS abundance are usually up-regulated together, more recent reports have shown that mucoidy can be regulated independently of CPS abundance.^4,5,9,42^ Our data agree with these reports as we observed that importable sugars all suppressed mucoidy, but some increased CPS abundance (*e.g.*, Xyl, GalNAc), some decreased CPS abundance (*e.g.*, Rha), and some did not affect CPS abundance (*e.g.*, Gal, Glc) (**Fig. 1C**).^5,6,20,22^ The independent effect of sugars on CPS abundance was especially striking with Gal and Glc as sole carbon sources, where the abundance increased by nearly two-fold, despite suppressing mucoidy over two-fold (**Sup. Fig. 2A-B**). Overall, these effects on CPS abundance did not significantly correlate with mucoidy (**Sup. Fig. 2C**). Thus, while sugars can have a variety of effects on CPS abundance, they broadly and independently suppress *K. pneumoniae* mucoidy.

RmpD is a key regulator of mucoidy in hypervirulent *K. pneumoniae*. The *rmp* operon, which encodes *rmpD*, is transcriptionally responsive to diverse environmental cues.^5,7,10–12^ For example, the arginine-responsive ArgR and iron-responsive Fur regulators have been shown to directly activate *rmpADC* promoter.^7,39^ In this study, we present that sugar-suppressed mucoidy is also mediated through transcriptional *rmpADC* regulation. Gal supplementation markedly reduced *rmpADC* promoter activity and relative transcript levels of all three genes within the operon (**Fig. 3A-B**). More broadly, the P*_rmp_* activation was consistently reduced across all mucoidy-suppressing sugars (**Fig. 4A**). Thus, transcriptional regulation of *rmpADC* could be a general response to sugar availability. Although Gal significantly reduced *rmpC* expression, it caused only a moderate reduction in CPS abundance. However, not all sugars had reduced CPS abundance despite a reduction in overall *rmpADC* promoter activation (**Fig. 1C** and **4A**). We previously observed a similar incongruity between CPS abundance and *rmpC* expression.^7^ CPS biosynthesis is regulated at multiple layers, and the effect of one regulator could be compensated for by another transcriptional regulator.^49^ Thus, a transcriptional change of one regulator, *rmpC*, might not translate to a proportional change in CPS abundance due to the activity of a hierarchically dominant regulator.†

To characterize the genetic basis of sugar-induced mucoidy suppression, we screened an ordered KPPR1 transposon library and validated 11 *K. pneumoniae* genes whose disruption overcame sugar-suppressed mucody (**Fig. 5A-B**). These genes are related to diverse cellular functions, including carbon-responsive global regulation (*cyaA*, *crp, csrD*), nitrogen regulation and sensing (*ntrB*, *ntrC*, *ptsN*), peptide transport (*sapA*, *sapB*), phosphate uptake (*pitA*), pentose phosphate pathway (*rpe*), and one gene of unknown function (*3906*). While several of these genes have been previously implicated in mucoidy or CPS modulation, they were not studied in the context of sugars as regulatory nutrient cues.^42,50^ Thus, it is important to note that some of the genes could be a general modulator of mucoidy independent of nutrient signal. For instance, *sapA* and *sapB* have been previously implicated in mucoidy and CPS regulation, but the underlying mechanism remains unresolved. Nevertheless, the dominance of carbon and nitrogen-associated genes in our screen re-affirms that metabolic status plays a major role in regulating *K. pneumoniae* mucoidy.^42^ Specifically, shifting the balance of carbon and nitrogen levels could drive *K. pneumoniae* to a hypo- or hyper-mucoid state. Since amino acids are positive modulators of mucoidy, we speculate that bacterial integration of sugars (carbon-rich) and amino acids (nitrogen-rich) availability in the host environment defines mucoidy levels. This is supported by the observation that sugar-suppressed mucoidy was concentration-dependent in M9+CAA+Gal, where mucoidy-activating arginine in CAA could oppose the mucoidy-suppressing effects of lower sugar concentrations (**Fig. 1B**). Thus, shifting metabolism in favor of either sugars or amino acids, could shift the cell to a hypo- or hyper-mucoid state. These findings collectively support the emerging view that *K. pneumoniae* mucoidy is controlled by complex and integrated regulatory networks.^42,49,51^ Rather than being regulated as a binary on/off trait, mucoidy is fine-tuned in response to multiple nutrient and environmental cues, providing niche-specific adaptations to the *K. pneumoniae* cell surface.

Although the transposon screen identified several candidates likely involved in sugar-suppressed mucoidy, we focused on *cyaA* and *crp* as CyaA activation and cAMP synthesis depend on sugar import. In Gram-negative bacteria, import of non-PTS or less-preferred sugars activates CyaA to synthesize the second messenger, cAMP.^52^ When complexed with CRP, cAMP-CRP functions as a global transcriptional regulator affecting several cellular processes, including virulence and fitness.^53–57^ Notably, CRP represses *K. pneumoniae* CPS biosynthesis by acting at multiple CPS biosynthesis promoters.^46,58^ It should be noted that not all sugars consistently changed CPS abundance despite a uniform decrease in intracellular cAMP (**Fig. 1C** and **7A**), suggesting that the regulatory influence of CRP may be context-dependent and modulated by other regulatory factors. Moreover, CRP has also been implicated in mucoidy suppression.^45,58^ Although the precise mechanism remains unclear, cAMP-CRP is proposed to suppress mucoidy by up- regulating the small RNA ArcZ, which inhibits the translation of transcripts associated with MlaA system.^45^ This system maintains outer-membrane asymmetry, which is essential for cell surface CPS retention. Therefore, ArcZ-dependent mucoidy regulation likely occurs independent of *rmpD* regulation and reflects altered CPS retention rather than chain length regulation. Moreover, the marked increase in *rmpADC* promoter activity observed in *cyaA* and *crp* mutants cannot be explained by ArcZ due to the post-transcriptional mode of regulation of ArcZ. Thus, the sugar-suppressed mucoidy reported here likely acts through a distinct mucoidy regulatory mechanism.

CRP-dependent promoters can be activated through several mechanisms depending on CRP binding site configuration and intracellular cAMP concentraiton.^59^ In *Escherichia coli*, for example, the *pck* promoter switches from activation to repression depending on cAMP-CRP levels.^60^ At low-[cAMP], CRP binds to the high-affinity binding site located upstream of the transcription start site and activates *pck*, but at high [cAMP], it binds to both high- and low-affinity binding sites causing an overall repression of *pck*.^60^ In our study, intracellular cAMP levels were elevated in the absence of sugar (M9+CAA), which correlated with increased mucoidy (**Fig. 7A** and **1A**). Conversely, reduced intracellular [cAMP] (M9+CAA+sugar) correlated with decreased mucoidy (**Fig. 7A** and **1A**). Thus, whether acting directly or indirectly at the *rmpADC* promoter, cAMP-CRP regulates *rmpADC* transcriptional expression in response to sugar import. However, the linear mucoidy differences in a high- and low-[cAMP] scenarios conflicts with *cyaA* and *crp* mutant phenotypes, both of which exhibit increased mucoidy in M9+CAA+Gal, despite undetectable intracellular [cAMP]. This paradox could be explained by considering one or more of the following regulatory scenarios. First, P*_rmp_* may contain multiple CRP-binding sites with varying affinities to cAMP-CRP, responding differently across cAMP-CRP concentrations, analogous to *E. coli pck* promoter. Second, CRP may concurrently regulate mucoidy directly through P*_rmp_* regulation and indirectly via other regulatory inputs. Third, CRP itself may be regulated by additional regulatory inputs. However, these proposed regulatory circuits remain speculative and must be validated with further studies. Even if cAMP-CRP does not bind the *rmp* promoter, it is likely that cAMP-CRP indirectly regulates the *rmp* locus in a [cAMP]-dependent manner.

*K. pneumoniae* infections often originate from gut colonization, where it primarily colonizes the small intestine followed by sustained persistence in the colon over time.^61^ Nutrient availability such as dietary simple carbohydrates are critical in the colonization and dissemination of *K. pneumoniae*.^62^ In fact, bacterial carbohydrate uptake and catabolic pathways, particularly galactose, fucose and xylose, are important for *K. pneumoniae* colonization in murine gut models.^11,63,64^ Based on this knowledge, we examined how Gal-suppressed mucoidy influences *K. pneumoniae* gut colonization. Within the colon, *K. pneumoniae* primarily localizes to the outer mucus layer while it can also invade the inner mucus layer adjacent to the colonic epithelium.^65^ Our data suggest that sugar-suppressed mucoidy may enhance the ability of *K. pneumoniae* to invade and interact with the intestinal mucin, as both *K. pneumoniae* pre-cultured in Gal and a constitutively non-mucoid strain significantly increased bacterial binding to gastric mucin **(Fig. 8A)**. Whether this modest difference in binding is sufficient to alter persistence in the gut *in vivo* remains to be seen. We also acknowledge that the type of mucin used in our study does not fully recapitulate the mucin diversity across the entire GI tract. To date, the most well-established function of mucoidy has been observed in limiting bacterial association with host cells *ex vivo*.^4,5,7,9^ A previous study demonstrated that *K. pneumoniae* can invade intestinal epithelial cells *in vitro*.^47^ Consistent with this, Gal-suppressed mucoidy increased the association of *K. pneumoniae* to intestinal epithelial cells, suggesting that sugar availability on host niches could dictate mucoidy state and facilitate bacterial interaction to host cells **(Fig 8B)**. Importantly, carbon sources are a limited resource in the gut environment for *K. pneumoniae,* and dietary carbohydrates can further influence the pool of available sugars. Therefore, dietary modulation of gut carbohydrates could also impact *K. pneumoniae* mucoidy during colonization. Since sugar import broadly suppresses mucoidy, the availability of even a subset of mucoidy-suppressing sugars could be sufficient to maintain a non-mucoid state in the gut. Thus, mucoidy is tightly aligned to the remarkable metabolic flexibility of *K. pneumoniae*.^66^

Future studies that map spatial and temporal regulation of mucoidy during different stages of colonization and dissemination will be critical for understanding how dynamic control of mucoidy contributes to *K. pneumoniae* infection. While our study focused on how sugars impact CPS-associated features, it is likely that sugar import and as a result, changes in cAMP-CRP alters other bacterial features important in the context of infection.^24^ Thus, sugars may serve as a broad nutrient cue that shape bacterial adaptation beyond cell surface polysaccharides. Building on this, our findings also raise intriguing questions about how *K. pneumoniae* integrates multiple and sometimes opposing nutrient cues. The concentration-dependent effects observed here indicate a potential competition between mucoidy-enhancing and mucoidy-suppressing cues in complex nutrient environments. How *K. pneumoniae* prioritizes between the opposing signals and what determines the dominance of one regulatory pathway over another remains to be studied. Moreover, we did not test all sugars at varying concentrations. Although high sugar concentrations uniformly suppressed mucoidy, whether individual sugars differ in mucoidy-regulating potency at lower concentrations could provide insight into hierarchical effects of individual sugars on suppressing mucoidy. At the mechanistic level, further studies are needed to assess regulation of *rmpADC* beyond the transcriptional level, as *rmpADC* promoter activity did not always correspond to mucoidy phenotype, as was observed in *cyaA* and *crp* mutants. Although the promoter activation in M9+CAA was similar between WT and the mutants, mucoidy was significantly lower in *cyaA* and *crp* than WT (**Fig 6A** and **6D**). This incongruity between *rmp* promoter activity and the mucoid phenotype suggests post-transcriptional regulation of *rmp* transcripts. CsrA is a post-transcriptional global regulator responsive to carbon availability and acid end-product accumulation. In *E. coli*, it is known to regulate bacterial motility, biofilm formation and metabolism by binding to transcripts and blocking ribosome binding.^67,68^ CsrA activity in *E. coli* is attenuated by the expression of small regulatory RNAs, *csrB* and *csrC,* which in turn are inhibited by CsrD.^67,69^ Notably, *csrD* is a hypermucoid transposon hit in our screen here and in other mucoidy screens.^42,50^ Thus, it is possible that post-transcriptional discrepancies between P*_rmp_* activity and mucoidy in the *cyaA* and *crp* could be due to CsrA activity. Interestingly, the cAMP-CRP and Csr cascades cross-regulate expression of one another in other Gram-negative enteric bacteria.^70–72^ Although it remains to be studied, these observations allude to a potential role for CRP and CsrA in maintaining mucoidy homeostasis in *K. pneumoniae* by acting at different stages of gene expression. Overall, these outstanding questions highlight the need to explore the association of nutrient fluctuations in the gut with the transition of *K. pneumoniae* from colonizer to invader state.

Altogether, we present a model in which cAMP-CRP-dependent regulatory processes broadly fine-tune *K. pneumoniae* mucoidy, thereby influencing key stages of *K. pneumoniae* infection, such as intestinal colonization **(Fig. 9)**. Information about sugar import is relayed through changes in intracellular cAMP-CRP levels. Specifically, intracellular cAMP-CRP is reduced following sugar import. Our results support that cAMP-CRP directly or indirectly modulates the *rmpADC* promoter, leading to altered *rmpD* transcription. RmpD interaction with Wzc then regulates CPS chain length uniformity and mucoidy. These sugar-driven changes in mucoidy alter bacterial binding to mucin and host epithelial cells, such as those in the GI tract and may facilitate gut colonization.

**Figure 9.**
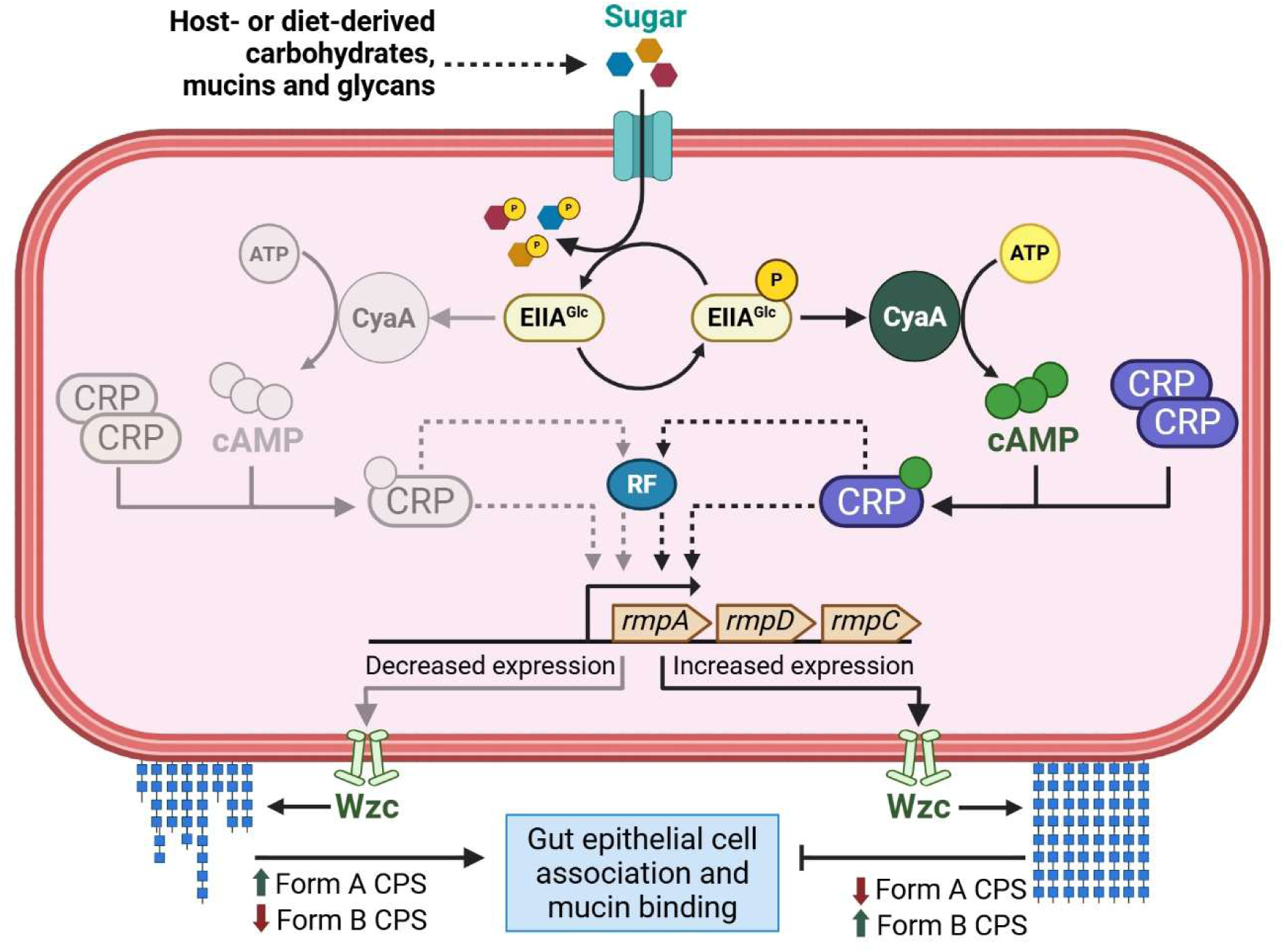
Model of sugar-suppressed mucoidy in hypervirulent *K. pneumoniae*. *K. pneumoniae* encounters ecological niches, such as the human gut, where it can import both mucoidy-inducing amino acids and mucoidy-suppressing sugars. In the absence of sugars, EIIA^Glc^ remains phosphorylated, which activates CyaA cAMP synthesis. Sugars import dephosphorylates EIIA^Glc^, which in an unphosphorylated state, is unable to activate the CyaA adenylate cyclase. cAMP forms a complex with the CRP transcription factor to regulate gene expression. The cAMP-CRP complex directly or indirectly via a yet-to-be identified regulatory factor (RF) modulates expression of the mucoidy regulator *rmpD*. Overall, effect of sugar import on lowering intracellular cAMP levels also reduces *rmpADC* expression. Reduced cellular levels of RmpD would diminish CPS chain length modulation via Wzc interactions, resulting in less uniform CPS chains on the bacterial cell surface (more Form A and less Form B CPS). Decreased CPS chain length uniformity is associated with suppressed mucoidy which promotes enhanced bacterial binding to gut mucin and intestinal epithelial cells. “RF” indicates a hypothetical regulatory factor influenced by cAMP-CRP. Dotted lines represent proposed interactions.

In summary, bacterial pathogens encounter nutritionally diverse environments that critically drives their adaptation for better fitness.^73^ Depending on the niche and stage of infection, the ability to integrate environmental cues and dynamically adjust metabolism, virulence and fitness determinants may be essential for bacterial persistence and successful infection. Our findings identify sugars as key nutrient signals that modulate *K. pneumoniae* cell surface properties through cAMP-CRP-dependent regulation of mucoidy and CPS chain length. While mucoidy provides protection against phagocytosis by blocking association to macrophages, it can simultaneously improve bacterial adherence to epithelial surfaces such as those of the gut, where attachment is likely important for persistence and dissemination. In this context, sugar-suppressed mucoidy may represent an adaptive response to host environments, where shifting nutrient profiles influence the bacterial metabolic state to promote colonization or invasive behaviors.

## 4. MATERIALS AND METHODS

### Bacterial strains and culture conditions

All bacterial strains and plasmids used in the study are detailed in **Supplementary Table S1**. Bacterial strains were cultured in either lysogeny broth (LB) (5 g/L yeast extract, 10 g/L tryptone, 0.5 g/L NaCl) or low-iron minimal medium (6 g/L disodium hydrogen phosphate, 3 g/L monopotassium phosphate, 0.5 g/L sodium chloride, 1 g/L ammonium chloride, 14.7 g/L calcium chloride, 120.3 g/L magnesium sulfate) with different carbon sources as described in the text, and incubated aerobically at 37°C and 200 rpm shaking, unless otherwise noted. Solid culture medium was prepared by adding 20 g/L bacto-agar to LB. When appropriate, antibiotics were added at the following concentrations: kanamycin (25 µg/mL), chloramphenicol (20 µg/mL *E. coli* or 80 µg/mL *K. pneumoniae*), gentamicin (10 µg/mL or 15 µg/mL).

### Sedimentation assay

Mucoviscosity (mucoidy) was quantified by using a low-speed centrifugation assay as previously described.^41,74^ Briefly, 1 mL of 0.5 OD_600_ equivalent bacterial overnight culture in 2 mL microcentrifuge tube was pelleted at 1000 *x g* for 5 min. The OD_600_ of the 900 µL supernatant was then quantified and presented as ‘supernatant OD_600_ per 0.5 OD_600_ bacterial culture’.

### Uronic acid quantification

Capsular polysaccharide content was measured by quantifying uronic acid content as previously described.^41,75^ In brief, capsular polysaccharide was purified by ethanol precipitation from 250 µL of overnight culture. 50 µL 1% Zwittergent 3-14 was added to overnight culture (1:5 ratio) to purify total capsular polysaccharide while 50 µL of ultra-pure water was added to purify cell-free EPS. The mixture was precipitated at 17,000 *x g* for 5 mins and the polysaccharide was collected by precipitating 100 µL of the uppermost supernatant in 400 µL ice-cold ethanol. Then, uronic acid content in the purified polysaccharide was quantified by using a colorimetric assay as previously described.^41^ Cell-associated CPS uronic acid level was deduced by subtracting the uronic acid levels in cell-free EPS from the total capsular polysaccharide.

### Molecular cloning, transformation and sequencing

Oligonucleotides used for cloning in this study are listed in **Supplementary Table S2**. *K. pneumoniae* isogenic knockout mutants were constructed using the λ Red recombineering adapted to *K. pneumoniae*.^42,76,77^ Fluorescent reporter plasmids were constructed by PCR amplification of the plasmid backbone (pBBR1MCS-5) and specific gene fragments (promoter and fluorescent protein) followed by gel-purification (Monarch, NEB), and assembled using NEBuilder HiFi DNA Assembly Mix (NEB) for 1 hour at 50°C. Similarly, *cyaA* and *crp* complementation vectors were constructed by ligating PCR-amplified and gel-purified vector backbone (pACYC184) and the respective gene fragments along with their native promoter. The resulting ligation production was then transformed into TOP10 *E. coli* through heat-shock and confirmed via sequencing. Electroporation of the final plasmid into *K. pneumoniae* strains was performed as previously described.^77^

Plasmid constructs were validated using whole plasmid sequencing (Plasmidsaurus and Eurofins Genomics). Select transposon mutants and all gene deletion mutants were validated using whole genome sequencing (SeqCoast Genomics).

### Growth curves

Growth curves were generated as previously described.^78^ In brief, overnight cultures were setup in the respective growth media and incubated aerobically at 37°C and 200 rpm. Then, a 1:1000 subculture of the overnight culture was prepared in 3 mL of the respective growth medium. One hundred µL of 0.001 OD_600_ equivalent subculture was then transferred to a sterile 96-well plate. Additionally, 100 µL of sterile water was added to the outermost wells of the plate to prevent evaporation of the samples. Then, OD_600_ was measured every 30 min for 16 hours at 37°C and continuous orbital shaking (282 rpm) using a microplate reader (EPOCH2-SN, Agilent). For growth media that were non-permissive to the bacterial growth, overnight culture and the subsequent subculture were prepared using M9 1% CAA.

### Fluorescent reporter assay

Intracellular cAMP level and *rmp* promoter (P*_rmp_*) activity were measured using one of the two fluorescent reporter constructs (**Table S1**). These reporter constructs were modified from a published fluorescent reporter vector.^79^ For experiments requiring measurement of P*_rmp_* only (**Fig. 3A** and **4C**), a single reporter construct expressing P*_rmp_*-*dasherGFP* was used.^7^ For all other experiments requiring measurement of both P*_rmp_* and cAMP, a double reporter plasmid expressing P*_rmp_*-*dasherGFP* and *lacP1*-*mScarlet-I* was used. In all cases, 1 mL of 0.5 OD_600_ equivalent culture was centrifuged at 21,000 *x g* for 10 min to remove culture medium. Following the centrifugation, the supernatant was completely removed, and the pellet was resuspended in 1 mL 1× PBS. Then, 300 µL of the resuspended cells were transferred to a clear-bottom, black 96-well plate. DasherGFP fluorescence intensity was measured using a microplate reader (Synergy HTX, Agilent) at excitation and emission wavelengths of 485 and 528 nm, respectively. Similarly, mScarlet-I fluorescence intensity was measured at excitation and emission wavelengths of 560 and 620 nm, respectively. The fluorescence intensity was normalized to OD_600_ and reported as an arbitrary fluorescence unit per OD_600_ (AFU/OD_600_).

### Transposon screen in sugar-supplemented growth medium

An ordered KPPR1 transposon library was used to screen for transposon mutants with increased mucoidy in galactose-supplemented medium.^42^ The screening and plate-based sedimentation assay were performed as previously described.^42^ Briefly, microplates containing the ordered library (a total of 3,733 mutants) were thawed at room temperature and replicated into 100 µL of M9 1% CAA + 80 mM gal. The replicated plates were then wrapped with a plastic wrap and incubated at 37°C with 200 rpm shaking for 18–20 h. Following incubation, the plates were vortexed at low speed for 30 seconds and measured for OD_600_ and centrifuged at 1000 *x g* for 5 min. Then, the upper 50 µL was transferred to a new microplate and the OD_600_ was measured. Transposon mutants with mucoidy (supernatant OD_600_ per culture OD_600_) greater than two times the standard deviation of the plate mean were considered hits of the primary screen. The primary hits were then arrayed in a new microplate for secondary screen. The newly arrayed plates contained each mutant in three different wells as a biological triplicate. The secondary screen was performed using a plate-based sedimentation assay and the hits were identified following an inclusion criterion as described for the primary screen. For validation, KPPR1 WT and the secondary screen hits were cultured in M9+CAA+Gal, quantified for mucoidy using tube-based sedimentation assay and considered a valid hit if its mucoidy was significantly higher than that of WT in M9+CAA+Gal.

### RNA isolation and quantitative RT-PCR

RNA isolation and q-RT-PCR were performed as previously described.^5,7^ Bacterial strain was cultured in 3 mL of low-iron M9 minimal media supplemented with 1% CAA with or without 80 mM galactose and incubated overnight aerobically at 37°C with 200 rpm shaking. Next day, overnight cultures were sub-cultured 1:100 in the respective medium and incubated aerobically at 37°C until the OD_600_ reached 0.4–0.5. Then, approximately 1 x 10^9^ colony forming units (CFUs) were mixed with RNAProtect (Qiagen) at 1:2 (sample:RNAProtect) ratio, incubated at room temperature for 5 min and the cells were collected by centrifugation at 5000 *x g* for 10 min. Then, pelleted cells were lysed with 100 µL lysozyme (15 mg/mL in TE buffer) and treated with 10 µL proteinase K (20 mg/mL in TE buffer). RNA was isolated and purified using RNeasy mini-prep kit (Epoch) following manufacturer’s instruction and eluted with 30 µL RNase-free water and stored at -20°C until downstream processing. Genomic DNA was removed from the RNA preparations using ezDNase (ThermoFisher). cDNA was synthesized from the purified RNA using SuperScript IV First Strand Synthesis System (Invitrogen) following the manufacturer’s directions. The resulting cDNA was diluted 1:50 in water and used as template for quantitative real-time PCR (qRT-PCR) with SYBRGreen PowerUp reagent (Invitrogen) in QuantStudio 3 PCR system (Applied Biosystem). Primers used to amplify *rmpA*, *rmpD*, *rmpC* and *gap2* (internal control) genes are listed in **Supplementary Table 2**.^5^ The relative fold change of the genes was calculated using the comparative threshold cycle (C_T_) method.^80^

### CPS chain length visualization

CPS chain lengths were visualized using as previously described.^5,41^ Cell-attached CPS were purified from bacteria equivalent to 1.5 OD_600_. Cells were pelleted at 21,000 × *g* for 15 min to remove cell-free EPS. The bacterial pellet was resuspended in 1 mL of PBS and centrifuged at 21,000 × *g* for 15 min. All but 250 µL of the supernatant was removed, mixed with 50 µL 3-14 1% Zwittergent and incubated at 50°C for 20 min. The incubated mixture was then centrifuged at 21,000 × *g* for 5 min to collect the upper 100 µL of the supernatant. The supernatant was added to a tube containing 400 µL ice-cold ethanol to precipitate polysaccharide. Purified cell-associated CPS was mixed with 4 × SDS gel loading dye (3:1, sample:dye) and resolved using 4-15% TGX stain-free precast gel (Bio-Rad). The electrophoresis was performed for 4.5 hours at 300V on ice at 4°C. After electrophoresis, the gel was stained with 0.1% alcian blue (0.1% wt/vol ThermoFisher Alcian Blue 8 Gx in stain base solution) for 1 h. It was then stained with Pierce Silver Stain Kit (ThermoFisher) following the manufacturer’s instructions.

### CPS chain length diversity quantification

The chain length diversity was quantified as previously described.^7^ Stained CPS gels were quantified using ImageJ V1.54r for Windows. Gel area of equal size was selected for each lane including an empty lane for background measurement and plotted using one-dimensional electrophoretic gel analysis feature. The one-dimensional gel profile was then plotted, and different CPS forms and baseline were defined in the lane profile. The area of each peak corresponding to a distinct CPS form was quantified. Measurements from an empty control lane were subtracted from each sample lane to adjust for background. The quantification values presented in the figures are log_2_ fold change relative to wildtype. Comparisons were made between samples and control loaded from the same gel to account for gel-to-gel variability.

### Mucin binding assay

Bacterial mucin binding assay was performed as previously described.^81^ A Nunc-Immuno^TM^ MaxiSorp^TM^ 96-well plate (Milipore Sigma; Cat No. M9410-1CS) was coated with 300 µL of 0.5 mg/mL crude porcine gastric mucin (PGM) in 0.1 M acetate buffer (pH 5.0). The coated plate was spun at 250 *x g* for 3 min and incubated at 37°C for 24 hours. Bacterial strains were cultured in 3 mL of low-iron M9 minimal media supplemented with 1% CAA with or without 80 mM galactose and incubated aerobically at 37°C with 200 rpm shaking for 16-18 hours. Approximately 10^5^ bacterial cells were added per well in triplicate, spun at 250 *x g* for 3 minutes to facilitate binding, and incubated statically at 37°C for 1 hour. After incubation, the wells were washed 15 times with 300 µL sterile PBS and treated with 200 µL 0.5% Triton-X100 in PBS for 30 minutes to release bound bacteria. Input and output bacterial cells were serially diluted and plated on LB agar for enumerating CFU.

### Epithelial cell association assay

Immortalized intestinal epithelial cells derived from a colorectal adenocarcinoma patient (Caco2, Sigma 86010202-1VL) were maintained in DMEM medium with L-glutamine, 4.5 g/L glucose and sodium pyruvate (Corning) supplemented with 20% heat-inactivated fetal bovine serum (Corning), 1% non-essential amino acids (Fisher Sci), 100 U/mL penicillin, and 100 µg/mL streptomycin in an atmosphere of 5% CO_2_. Bacterial strains were cultured in 3 mL of low-iron M9 minimal media supplemented with 1% CAA with or without 80 mM galactose and incubated aerobically at 37°C with 200 rpm shaking for 16-18 hours. Caco-2 cells in a tissue culture-treated petri dish were seeded to a density of 75,000 cells/mL per well to a tissue culture-treated 24-well plate. The seeded cells were incubated for 24-48 hours to achieve a 90% confluent monolayer. Confluent Caco-2 cells in 24-well tissue culture dishes were washed with 1 mL of DPBS, then 1 mL bacteria (MOI 10) in additive-free DMEM was added to each well. Samples were spun at 500 rpm (54 × *g*) for 5 min, then incubated at 37°C, 5% CO_2_ for 1.5 and 3 hours. After incubation, samples were washed three times with PBS followed by lysis with 1 mL of 0.2% Triton-X100 in PBS for up to 10 min. Input and cell-associated bacterial counts were determined by serial dilution and CFU enumeration on LB agar.

### Statistics

All replicates represent biological replicates and were performed at least three times independently unless otherwise noted. Statistical analyses were computed in Graphpad Prism 10 (V 10.4.1; GraphPad Software, LLC). For experiments comparing more than two groups on two independent variables, significance was calculated using two-way ANOVA with Dunnett’s post-hoc test or Šídák correction. Meanwhile, one-way ANOVA with either Dunnett’s post-hoc test or Šídák correction was applied to calculate significance between more than two groups on one independent variable. Statistical differences between two groups with independent variables were calculated using student’s t-test. In all instances, results were considered statistically significant if the *P*-value was less than or equal to 0.05. Unless otherwise noted, the graphs represent an average of replicates with standard error of mean.

## ACKNOWLEDGEMENTS

Research reported in this publication was supported by the University of Toledo, University of Pittsburgh, 23CDA1056712 (L.A.M.) and 24PRE1197026 (S.K.) from the American Heart Association, and K22 AI145849 (L.A.M.) and R35 GM150588 (L.A.M.) from the National Institutes of Health. This content is solely the responsibility of the authors and does not necessarily represent the official views of the funding agencies.

We thank Drs. Matthew Parsek and Xuhui Zheng for the pBBR1 reporter plasmid backbone. We thank members of the Mike Lab (Brooke E. Ryan, Emily L. Kinney, Grace E. Shepard, Zachary J. Resko, Lindsey R. Krzeminski, Krista Pettee, Jolie G. Lagger), Department of Medical Microbiology and Immunology at the University of Toledo (Drs. R. Mark Wooten, Robert Blumenthal, Jason Huntley, Peter Andreana) and Program in Microbiology and Immunology/Division of Infectious Diseases at the University of Pittsburgh (Drs. William H. DePas, Matthew Culyba, Anthony Richardson and Patricia Grace) for sharing technical resources, and valuable guidance and feedback.

---

**Supplementary Figure 1.**
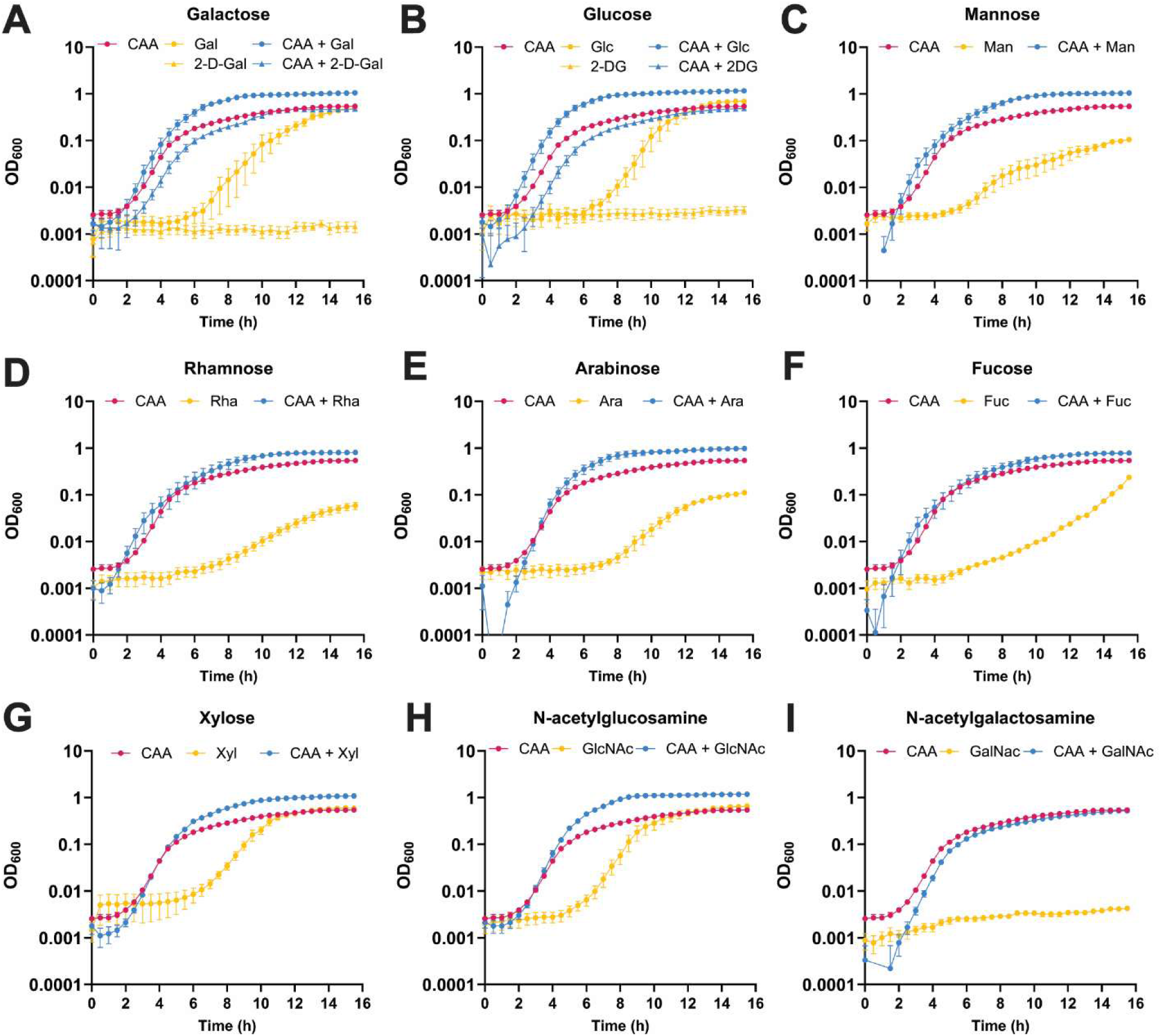
*K. pneumoniae* grows on most individual sugars, which increases when combined with casamino acids. KPPR1 was cultured in M9+CAA (1%), M9+sugar (80 mM) and M9+CAA+sugar. After inoculating KPPR1 in each growth medium, OD_600_ was measured every 30 minutes for 16 hours at 37°C. Data presented are the mean, and error bars represent the standard error of the mean. Experiments were performed 3 independent times, in triplicate.

**Supplementary Figure 2.**
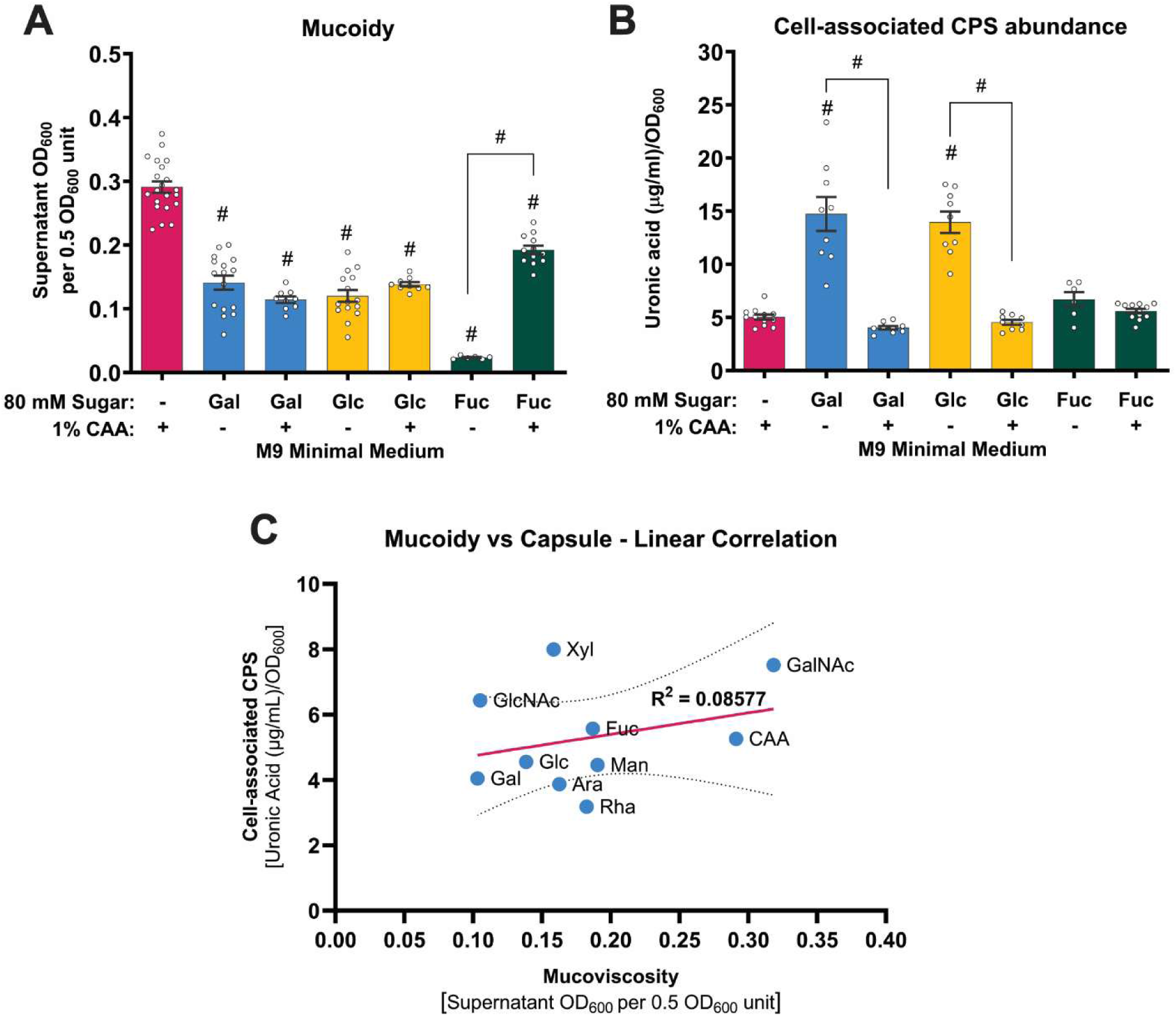
Sugars suppress *K. pneumoniae* mucoidy independent of CPS abundance. **(A** and **B)** KPPR1 was cultured in M9+CAA, M9+sugar and M9+CAA+sugar. **(A)** Mucoidy was determined by quantifying the supernatant OD_600_ after sedimenting 0.5 OD_600_ unit of culture at 1,000 x *g* for 5 mins and **(B)** uronic acid abundance was quantified for total CPS and supernatant CPS and normalized to OD_600_. Cell-associated CPS abundance was calculated by subtracting supernatant CPS from the total CPS content. **(C)** Mucoidy and cell-associated CPS abundance presented in Fig. 1A and 1C were analyzed for correlation using simple linear regression. **(A-B)** Data presented are the mean, and error bars represent the standard error of the mean. Statistical significance was determined using one-way ANOVA with Šídák correction. Statistical significance was calculated by comparing sugar-supplemented condition to M9+CAA or between adjacent pairs of bars. # *p* ≤ 0.0001. Experiments were performed ≥3 independent times, in triplicate.

**Supplementary Figure 3.**
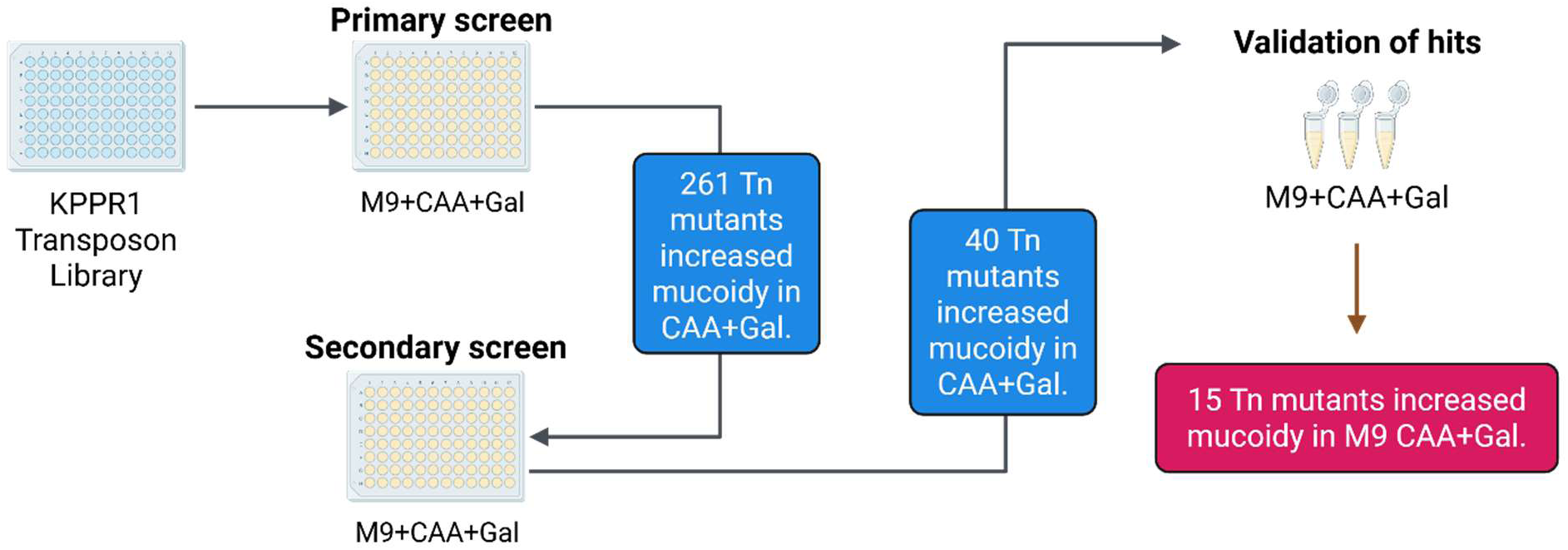
Graphical illustration of transposon screening workflow to identify mutants with increased mucoidy in galactose-supplemented medium. Primary and secondary screen were performed in M9+CAA+Gal using a sedimentation assay adapted to a 96-well plate format for high-throughput screening. Final validation of hits was performed by a tube-based sedimentation assay. Data are presented in Figure 5A. Mutants with mucoidy higher than two times the standard deviation of average mucoidy of a plate were identified as primary hits. Mutants with mucoidy significantly higher than WT in M9+CAA+Gal were confirmed as validated hits (*n = 15*).

**Supplementary Figure 4.**
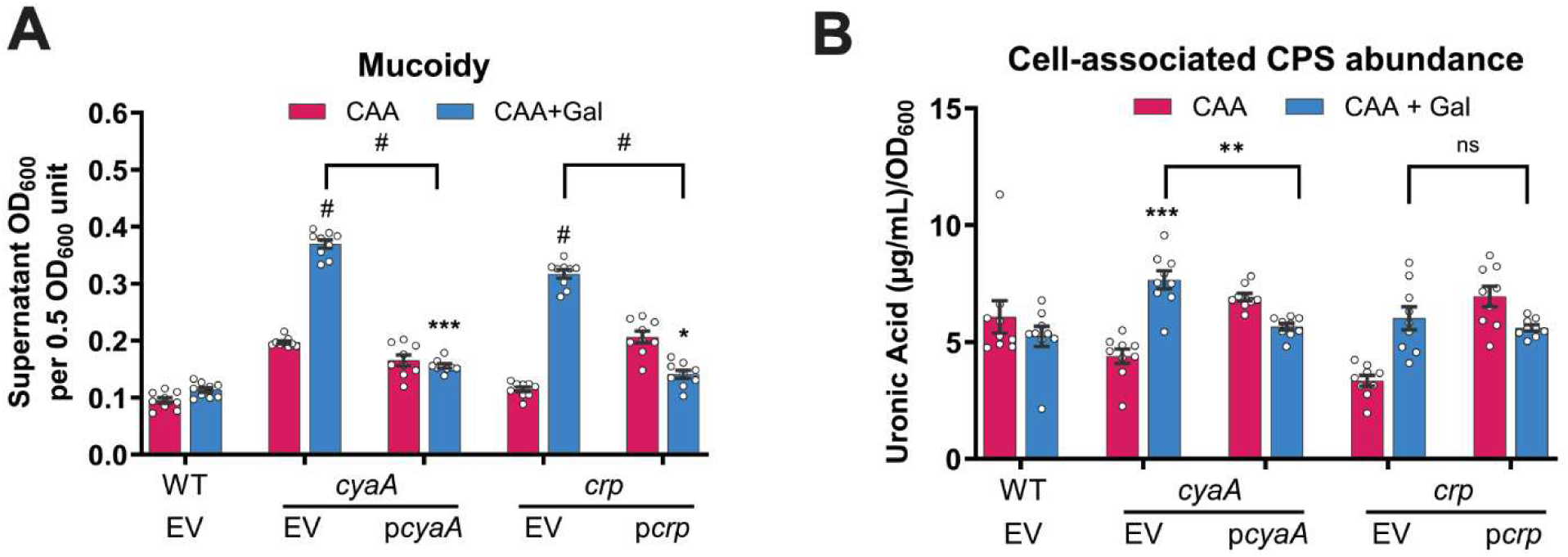
Complementation of *cyaA* and *crp* deletion strains. KPPR1 WT with empty vector (WT+EV), and *cyaA* and *crp* mutants with EV (*cyaA*+EV and *crp*+EV) or their respective complementation vectors (*cyaA*+p*cyaA* and *crp*+p*crp*) were cultured in M9+CAA±Gal. **(A)** Mucoidy was determined by quantifying the supernatant OD_600_ after centrifugation at 1,000 x *g* for 5 mins. **(B)** Cell-associated CPS was extracted and measured for uronic acid content. Data presented are the mean, and the error bars represent the standard error of the mean. Statistical significance was determined using two-way ANOVA with Šídák correction, by either comparing mutant-based strains in M9+CAA+Gal to WT in M9+CAA+Gal (*p*-value above each bar) or EV to the respective complementation vector (*p*-values above each connected line). * *p* ≤ 0.05; ** ≤ 0.01; *** *p* ≤ 0.001; # *p* ≤ 0.0001. All experiments were performed ≥3 independent times, in triplicate.

**Supplementary Figure 5.**
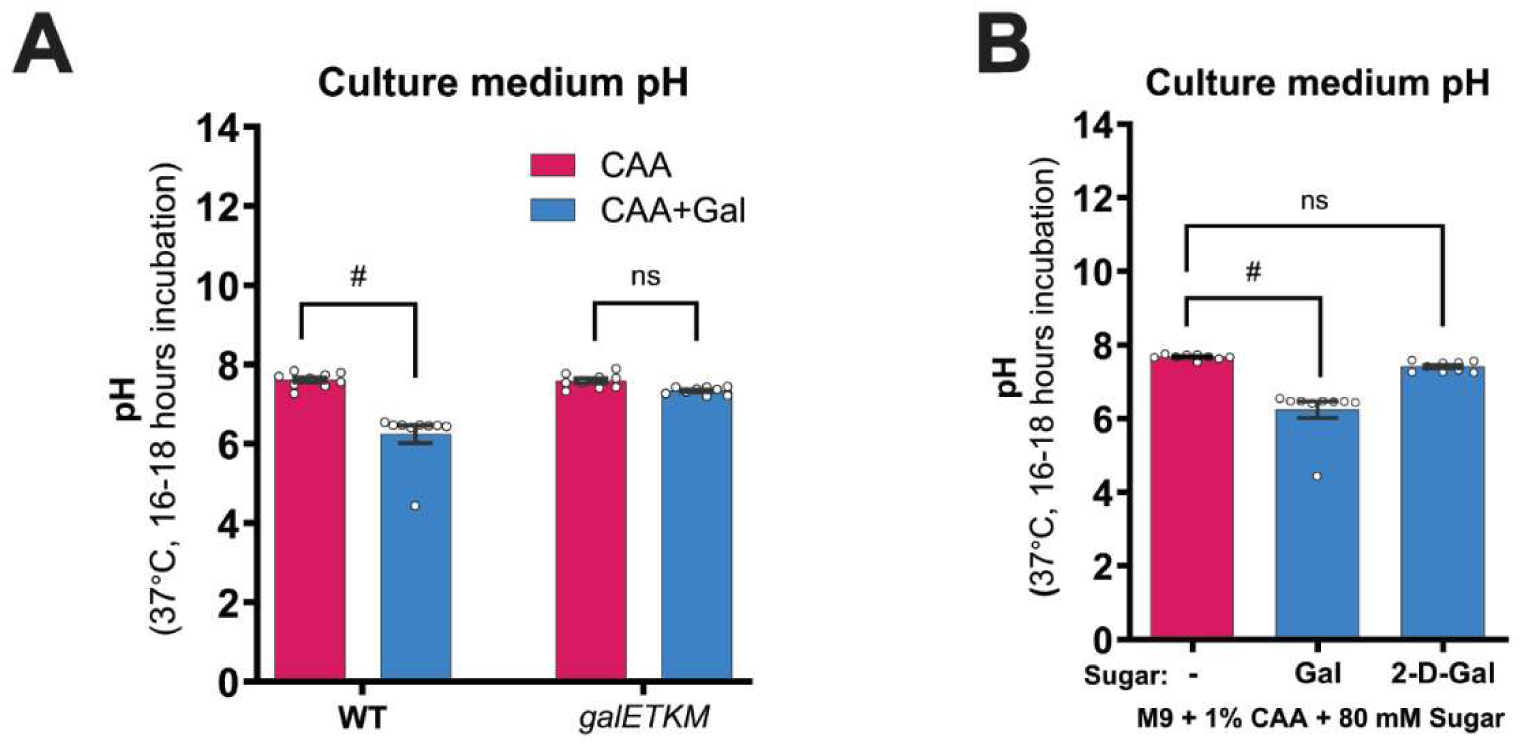
KPPR1 *galETKM* mutant and 2-deoxy-D-galactose do not decrease growth medium pH. **(A)** KPPR1 WT and *galETKM* mutant were cultured in M9+CAA±Gal. **(B)** KPPR1 WT was cultured in M9+CAA supplemented with Gal or the non-metabolizable analog, 2-D-Gal. For both **(A** and **B),** bacterial strains were cultured in their respective growth media at 37°C for 16-18 hours, then the culture medium pH was measured using a pH meter. Data presented are the mean, and error bars represent the standard error of the mean. Statistical significance was determined using **(A)** two-way ANOVA with Šídák correction or **(B)** one-way ANOVA with Dunnett’s post-hoc test. Statistical significance was calculated by comparing sugar-supplemented condition to M9+CAA. # *p* ≤ 0.0001; ns = non-significant. Experiments were performed 3 independent times, in triplicate.

